# Neuromodulation enables transient flexible control of motoneurons

**DOI:** 10.64898/2026.05.01.721852

**Authors:** Natalia T. Cónsul, Simon Avrillon, Mario Bräcklein, Juan A. Gallego, Dario Farina

**Affiliations:** Department of Bioengineering, Imperial College London, UK; Nantes Université, Laboratory “Movement, Interactions, Performance”, Nantes, France; Reality Labs at Meta, New York, NY, USA; Neuroscience of Disease Program, and Centre for Restorative Neurotechnology, Champalimaud Foundation, Lisbon, Portugal

## Abstract

A motoneuron pool is often regarded as a *rigid* controller because the largely shared synaptic input across motoneurons leads to strongly correlated activity. However, brief deviations from this correlated behavior have been observed even in some constrained tasks, raising the question of whether these results reflect limitations of the rigid view of motoneuron pool control. Here we show that they do not. We developed a biophysical model of a motoneuron pool receiving shared excitatory and inhibitory synaptic inputs that also included the motoneuron-specific effects of neuromodulation; model parameters were tuned based on large-scale motoneuron recordings in humans. Simulations showed that the intrinsic differences in how motoneurons respond to neuromodulation are both necessary and sufficient to transiently decorrelate pairs of motoneurons receiving a shared synaptic input. Crucially, such transient decorrelation is only observed when motoneurons have different sensitivity to neuromodulation, consistent with experimental observations during volitional control in humans. Our model also explains how participants can improve their ability to transiently decorrelate the activity of motoneurons innervating the same muscle by leveraging refined behavioral strategies that exploit the differential response of motoneurons to neuromodulation, rather than through physiological changes. These results identify that heterogeneous sensitivity to neuromodulation enables brief flexibility in the otherwise rigid control of motoneurons enforced by a shared synaptic input, and show how practice allows participants to exploit latent flexibility within otherwise rigid constraints.

## Introduction

Muscle force is controlled by populations of α-motoneurons in the spinal cord that innervate skeletal muscle fibers and drive their contraction. The motoneurons that innervate a single muscle, or a compartment of a multifunctional muscle, form a motoneuron pool. Motoneurons within a pool receive largely shared (excitatory) synaptic input^1,2^, which leads to orderly recruitment according to soma size as described by Henneman’s size principle^3^. This shared input also causes motoneurons in the same pool to exhibit highly correlated activity^4^, a behavior often referred to as “rigid control” of motoneurons.

Recent studies suggest that the nervous system may have more fined grained access to motoneurons innervating upper limb muscles than traditionally thought; in other words, there may be smaller “controllable units” than the entire population of motoneurons innervating a muscle. For example, in monkeys, intracortical microstimulation of primary motor cortex at different sites can consistently recruit the same two motoneurons in different order^5^, a finding anticipated by previous observations in baboons^6^. Deviations from classic rigid control also occur when comparing recruitment across different isometric tasks^5^, when a muscle changes its primary function^7,8^, or when the same task is being performed with different limb configurations that change muscle length^5^. Consistent with this increased control granularity, human participants can readily decorrelate the activity of motoneurons from the biceps muscle^9^. Therefore, the nervous system may have access to smaller functional units than the full motoneuron pool —at least for multifunctional muscles—, but whether each of these “submodules”^10^ can deviate from classic rigid control remains unclear.

The experiments by Bräcklein et al.^11^ provide insights into the potential flexibility of the control of these motoneuron submodules. These authors trained participants to decorrelate motoneurons innervating the same leg muscle (tibiliais anterior) using a cursor control task. Consistent with a classic rigid control view, none of the participants could change the recruitment order of a pair of motoneurons even after 14 days of practice. Yet, they became better at transiently decorrelating the motoneurons during derecruitment: with practice, participants learned to keep the putatively larger motoneuron active after the small one went quiet. This suggests that populations of motoneurons that are rigidly controlled can exhibit some transient flexibility during derecruitment that can be harnessed under voluntary control.

Here, we hypothesized that these deviations, which enable transient flexible control, arise from neuromodulatory inputs that alter motoneuron excitability even under a single shared input and without violating orderly recruitment (Fig. 1). In the case of spinal motoneurons, neuromodulatory input is provided by neurotransmitters such as serotonin and norepinephrine that are released from descending brainstem pathways^12,13^. Neuromodulation primarily affects motoneuron activity through persistent inward currents (PICs) that amplify synaptic inputs, promoting sustained firing after synaptic input cessation (hysteresis), and increasing discharge rates with repeated activation (facilitation)^14^. Crucially, these neuromodulatory effects are activity dependent: PICs are engaged only after motoneurons begin to fire^13^. As a result, recruitment order is largely preserved, whereas derecruitment can be strongly shaped by neuromodulation —a notion that is fully consistent with the results by Bräcklein et al.^11^, where transient decorrelation between motoneuron pairs only happened during derecruitment.

**Fig. 1.**
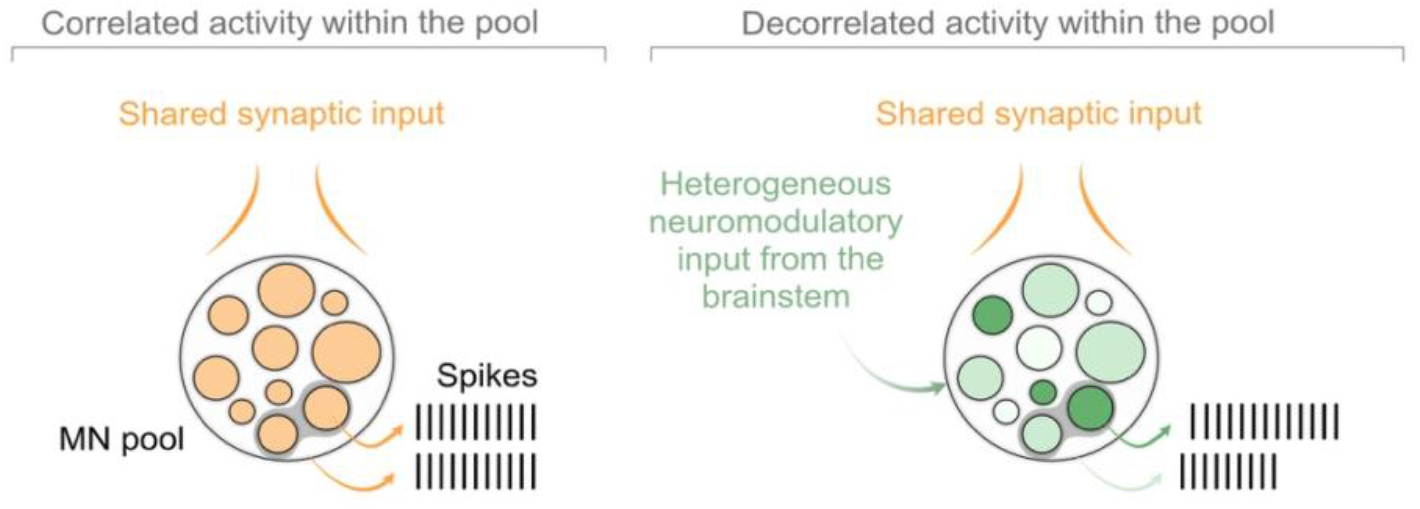
Hypothesis: descending heterogeneous neuromodulatory input from the brainstem decorrelates motoneuron activity. With a shared (excitatory) synaptic input, motoneurons within a pool fire in a highly correlated manner. Descending neuromodulatory input from the brainstem affects each motoneuron heterogeneously, and along with the unevenly distributed inhibitory synaptic input, leading to more decorrelated firing within the pool.

How can neuromodulation change the derecruitment order of spinal motoneurons under rigid control? Anatomical reconstructions have shown that serotonergic and noradrenergic inputs are densely but not uniformly distributed across motoneuron dendrites^15^, implying heterogeneous neuromodulatory effects within a pool. Together with evidence that inhibitory inputs to motoneurons may also be unevenly distributed^16,17^, this suggests that motoneurons receiving the same shared (excitatory) synaptic input may respond differently to the same neuromodulatory input, producing the partly decorrelated activity during derecruitment observed experimentally^11^.

Here, we asked whether neuromodulatory inputs are *necessary* and *sufficient* to decorrelate motoneurons under rigid control, that is, when a motoneuron pool receives shared synaptic input producing correlated activity^1,2^. We developed a biophysical model incorporating key neuromodulatory processes, including hysteresis and facilitation^14^, and calibrated it to large-scale human motoneuron recordings. Using this model, we show that the intrinsic differences in sensitivity to neuromodulation across motoneurons, in combination with their heterogeneous response to inhibitory inputs, is sufficient to generate transient decorrelation among motoneurons during derecruitment. We further demonstrate that the improved ability to transiently decorrelate motoneurons with practice reported by Bräcklein et al.^11^ can be explained by small adjustments in their behavioral strategies that better exploited the differential sensitivity to neuromodulation across motoneurons; no changes in the structure and properties of neuromodulatory inputs are needed. Our findings thus demonstrate that heterogenous neuromodulatory input, along with the unevenly distributed inhibitory inputs^16,17^, can generate brief departures from the normally correlated activity of motoneurons, and that humans can learn to better exploit this transient flexibility with practice.

## Results

We built a biophysical model of a human motoneuron pool to understand the effects of neuromodulation on the coordination among motoneurons receiving shared (excitatory) synaptic input, and its potential effect to transiently deviate from a rigid control scheme. Our model included excitatory and inhibitory synaptic inputs together with intrinsic channel dynamics to reproduce physiological variability. Model parameters were optimized based on experimental data from eight participants performing a force-tracking task^18^. Using this trained model, we simulated the online control experiments by Bräcklein et al.^11^ to address the hypotheses that neuromodulation is necessary and sufficient to transiently decorrelate the activity of biophysically distinct motoneurons during derecruitment, and that refined behavioral strategies can be learned to exploit this effect.

### Model architecture

We implemented a biophysical model of a motoneuron with two compartments, a soma and a dendrite, each with multiple conductances and channels (Fig. 2a). A motoneuron pool was then constructed by linearly interpolating the physiological parameters between the smallest and largest motoneurons, reproducing their natural dependence with size^19^. The number of motoneurons for each simulated pool was the same as those in the experimental recordings we used for tuning the model parameters^18^ (129 ± 44, mean ± s.d.; range, 62–214). Each motoneuron received shared excitatory and inhibitory synaptic inputs and was subject to the effects of neuromodulation; the three of them together were used to simulate the trains of motoneuron action potentials. These trains of action potentials were converted into force using a standard twitch-based^20^.

**Fig. 2.**
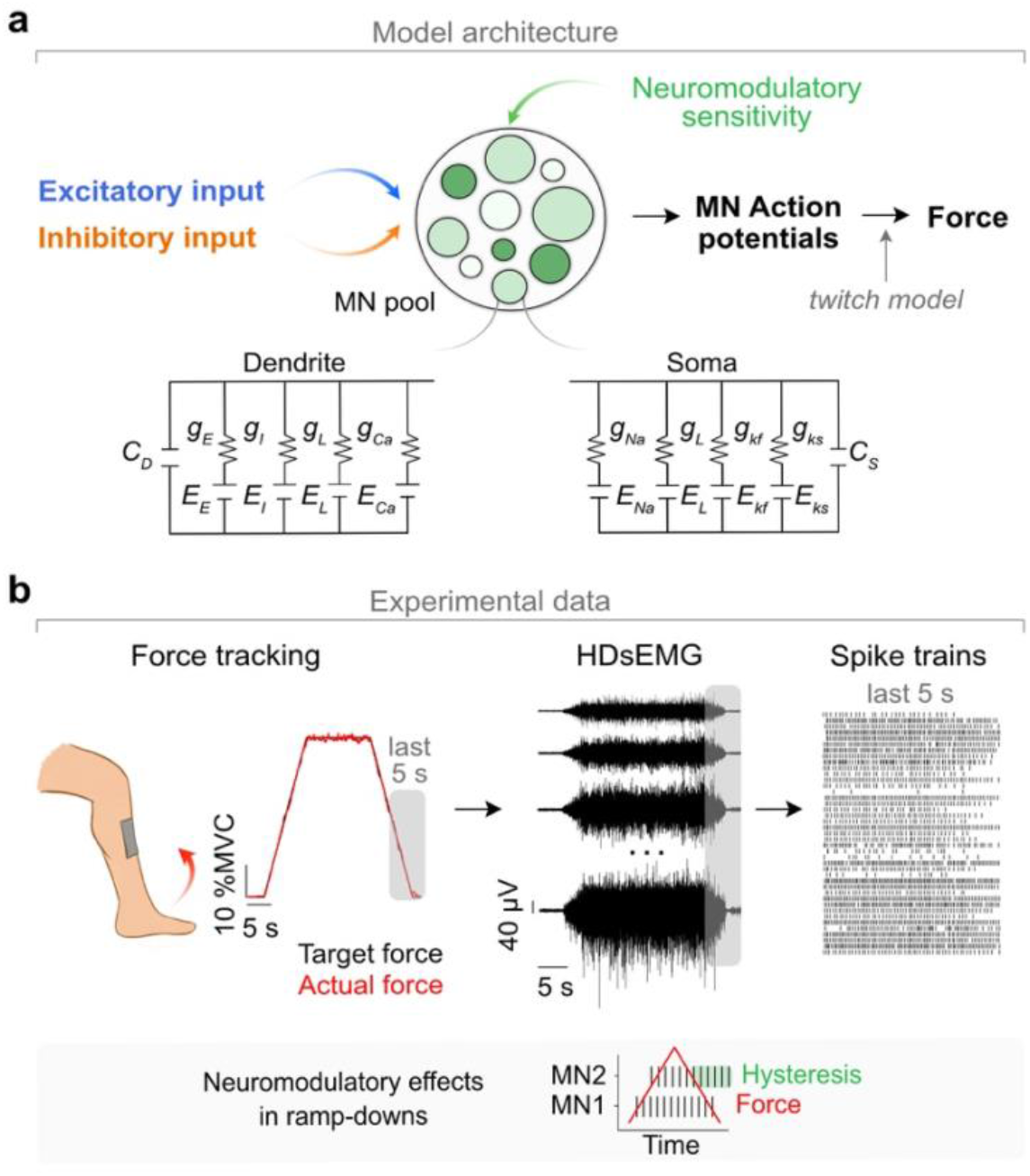
Model architecture and experimental recordings used for parameter tuning. **a:** Each motoneuron received shared excitatory and inhibitory synaptic inputs. Neuromodulatory effects emerged through the heterogeneous intrinsic sensitivity of calcium channels. The motoneuron model comprised two compartments, a dendrite and a soma. The dendrite included excitatory (E), inhibitory (I), and leak (L) conductances, as well as calcium (Ca) channels. The soma contained leak, sodium (Na), fast potassium (Kf), and slow potassium (Ks) channels (definitions in Methods). **b:** High-density surface electromyograms (HDsEMG; 256 channels) from the tibialis anterior muscle were collected as eight participants tracked trapezoidal force profiles by performing isometric contractions at multiple force levels (middle, four example EMG channels). Right: spiking of 181 motoneuron from a representative participant during the final 5 s of the ramp-down phase at 20% MVC, showing examples of hysteresis, as defined in the bottom schematic (sustained firing after input cessation, with derecruitment occurring at lower force levels than recruitment when motoneurons are ordered by recruitment threshold).

The neuromodulatory processes that we aimed to model could be potentially caused by either persistent sodium or calcium inward currents^14^. Calcium-mediated PICs exhibit slower dynamics and stronger hysteresis than their sodium counterparts^21,22^. Thus, we focused on calcium-mediated PICs since they were the most likely to mediate the experimentally-based effects during derecruitment^11^. Differences in intrinsic properties across model motoneurons lead to heterogeneous neuromodulatory effects across the pool, as observed in experiments ranging from decerebrated cats to behaving humans^4^.

### Parameter tuning and model validation

We tuned the model parameters based on the activity of populations of motoneurons from the tibialis anterior muscle —the same muscle by Bräcklein et al.^11^—, which we detected via decomposition of high-density surface electromyography recordings on eight participants^18^ (Fig. 2b). Motoneuron activity was detected across a broad range of muscle forces (10–80% of the maximum voluntary contraction, MVC), ensuring that the recordings included motoneurons of a broad range of sizes^3^. To fit the model parameters, we focused on recapitulating the persistent firing of motoneurons due to neuromodulation^14^ while capturing the interactions between inhibition and neuromodulation^13,23,24^, as well as replicating motoneuron facilitation during repeated contractions^14^. By building eight participant-specific models, we verified that our results generalized beyond a single set of parameters.

We first defined the relationship between inhibitory and excitatory synaptic inputs and the calcium voltages that determine how neuromodulation influences motoneuron activity. To elicit neuromodulatory effects and tune their calcium channel parameters, model motoneurons had to be recruited and derecruited; we thus applied a triangular excitatory synaptic input that recruited the full pool in order of motoneuron size. Inhibitory input was modelled as a constant input^25^, but we observed the same overall behavior in terms of recruitment and response to neuromodulation for all three commonly assumed relationships between excitatory and inhibitory synaptic inputs: constant, proportional, and inversely proportional^26^ (Fig. 3a, Suppl. Fig. 1a, 2). In the reminder of the paper, we used a constant relationship between excitatory and inhibitory synaptic inputs, except when indicated otherwise, including for controls.

**Fig. 3.**
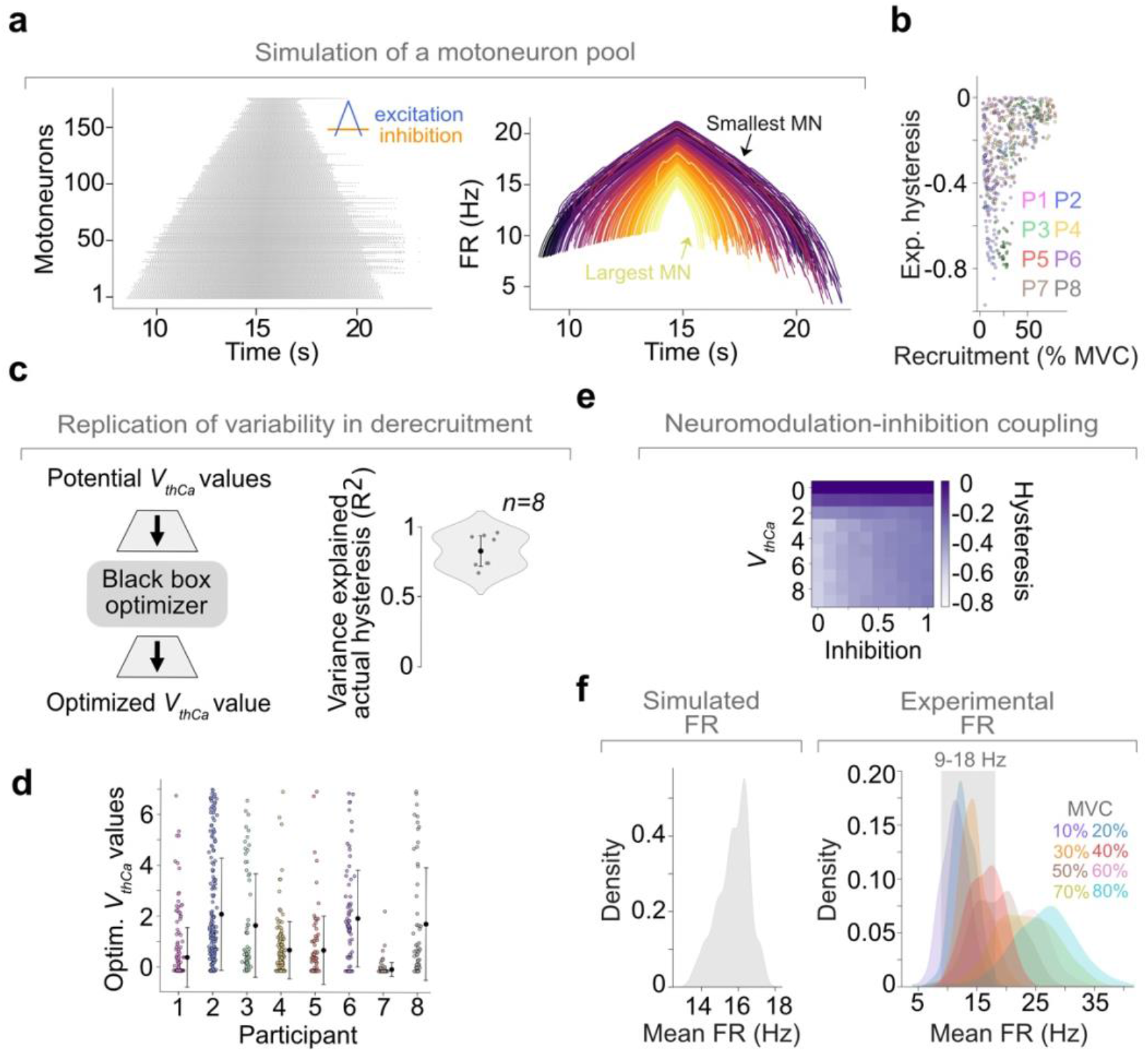
Parameter tuning and model validation. **a:** Example simulation of a motoneuron pool for a representative participant. Left: raster plot of the motoneuron pool (*n* = 181). Right: corresponding firing-rates (different colours) showing orderly recruitment and variable derecruitment times (note the mixing of colours). **b:** Experimental hysteresis for each participant (*n* = 8) varied systematically with recruitment threshold. Markers, motoneurons. **c:** Replication of the experimental variability in motoneuron derecruitment times. A black-box optimizer searched over potential *V*_*thCa*_ values to identify those that best reproduced the experimental hysteresis values for each participant, achieving high accuracy (R^2^). Markers, individual participants; the distribution includes all the motoneurons from all participants. **d:** Optimized *V*_*thCa*_ values showed substantial variability within and across participants. Markers, motoneurons; bars, s.d. of optimized *V*_*thCa*_ values for each participant. **e:** Our model recapitulated the experimental observation that hysteresis (colour) decreases with increasing inhibition amplitude. **f:** The firing rate distribution of simulated motoneurons (range, ~9–18 Hz) recapitulated the range of maximum overlap across experimental firing rate distributions at different contraction levels.

The parameters for our eight participant-specific models were fitted based on the experimentally observed effects of neuromodulation. These effects were quantified based on their hysteresis, defined as sustained firing after input cessation^14^, and calculated as the normalized ratio between the forces at derecruitment and recruitment, with more negative values indicating stronger effects (Fig. 3b). We used a black-box optimizer^27^ to tune the voltage threshold for calcium-channel deactivation (*V*_*thCa*_), achieving good fits to experiments values (mean R^2^ = 0.85; Fig. 3c,d). The optimized *V*_*thCa*_ values caused substantial hysteresis variability both across the motoneuron pool and across participants, consistent with experimental observations (Fig. 3b).

Next, we ensured that the model reproduced the expected interaction between inhibition and neuromodulation^13,23,24^ by which increasing inhibition should counteract the effects of neuromodulation and reduce the hysteresis. We built this relationship into our model by making inhibition inversely proportional to each motoneuron’s *V*_*thCa*_ value. Simulated hysteresis values were then computed across combinations of inhibition (range, 0–1; 0, no inhibition; 1, equal amplitude to excitation) and *V*_*thCa*_ (1–10) during a 30 s triangular contraction. As expected, higher *V*_*thCa*_ increased hysteresis, whereas stronger inhibition reduced it (Fig. 3e).

While the effects of neuromodulation during isolated muscle contractions mostly manifest as hysteresis, repetitive contractions engage the additional process of facilitation^14^, which we also built into our model. Under facilitation, motoneurons receiving stronger neuromodulatory input fire earlier and at higher rates after repeated contractions. We simulated this process as motoneurons with larger *V*_*thCa*_ undergoing a stronger activity-dependent shift in their calcium activation threshold after repeated contractions.

Finally, we verified that the model generated biologically realistic motoneuron pool behavior across a broad range of conditions. We implemented a proportional–derivative (PD) controller that modulated the shared excitatory synaptic input (which captured, among other factors, descending commands from the brain^28,29^), and used it to track diverse target force profiles^30–32^, including trapezoids, sinusoids, and chirp signals (Suppl. Fig. 1b). Under all these conditions, the firing rates of the simulated motoneuron pool fell within the experimentally observed range (~9–18 Hz), which we calculated as the range of maximum overlap across the firing rate distributions at the different force levels (Fig. 3f; this frequency range was biased towards the lower MVC levels due to the predominance of slow, low-threshold motoneurons that remain active across contractions, a behavior well captured by our model since it uses slow-type motoneurons^33,34^). Moreover, both during single and repetitive contractions, pairs of simulated motoneurons readily exhibited different behaviors, including deviations from correlated activity during derecruitment, which only emerged in pairs of motoneurons that were subject to different levels of neuromodulation (Suppl. Fig. 3). In sum, our model recapitulates all the necessary phenomena to test the hypothesis that neuromodulation enables transient flexible control of motoneurons.

### Neuromodulation is both necessary and sufficient to replicate the experimentally observed decorrelation between motoneuron pairs

Having showed that our model captures the variability within a motoneuron pool observed experimentally, we next asked whether neuromodulation could mediate the change in motoneuron derecruitment order observed during online cursor-control^11^. In this task, participants (*n* = 6) controlled the position of a cursor by modulating the activity of two motoneurons from the tibialis anterior muscle, which were selected for having similar recruitment thresholds to make the task sufficiently challenging. The activity of the lower-threshold motoneuron, motoneuron 1 (MN1), was mapped onto the position of the cursor along the horizontal axis, whereas that of the higher-threshold motoneuron (MN2) was mapped onto the position along the vertical axis. Although participants were given three targets (0º, 45º and 90º with respect to the horizontal, with the angles measured in the counterclockwise direction), here we focus exclusively on the 90º target, because this is the target for which derecruitment order had to be inverted. That is, to acquire the 90º target, participants had to sustain the activity of the higher threshold (i.e., larger) motoneuron (MN2) after the lower threshold motoneuron had stopped firing^11^ (MN1; “Transient flexibility” in Fig. 4a; note that participants could never activate MN2 without activating the smaller MN1^11^ to achieve full “Flexibility”).

**Fig. 4.**
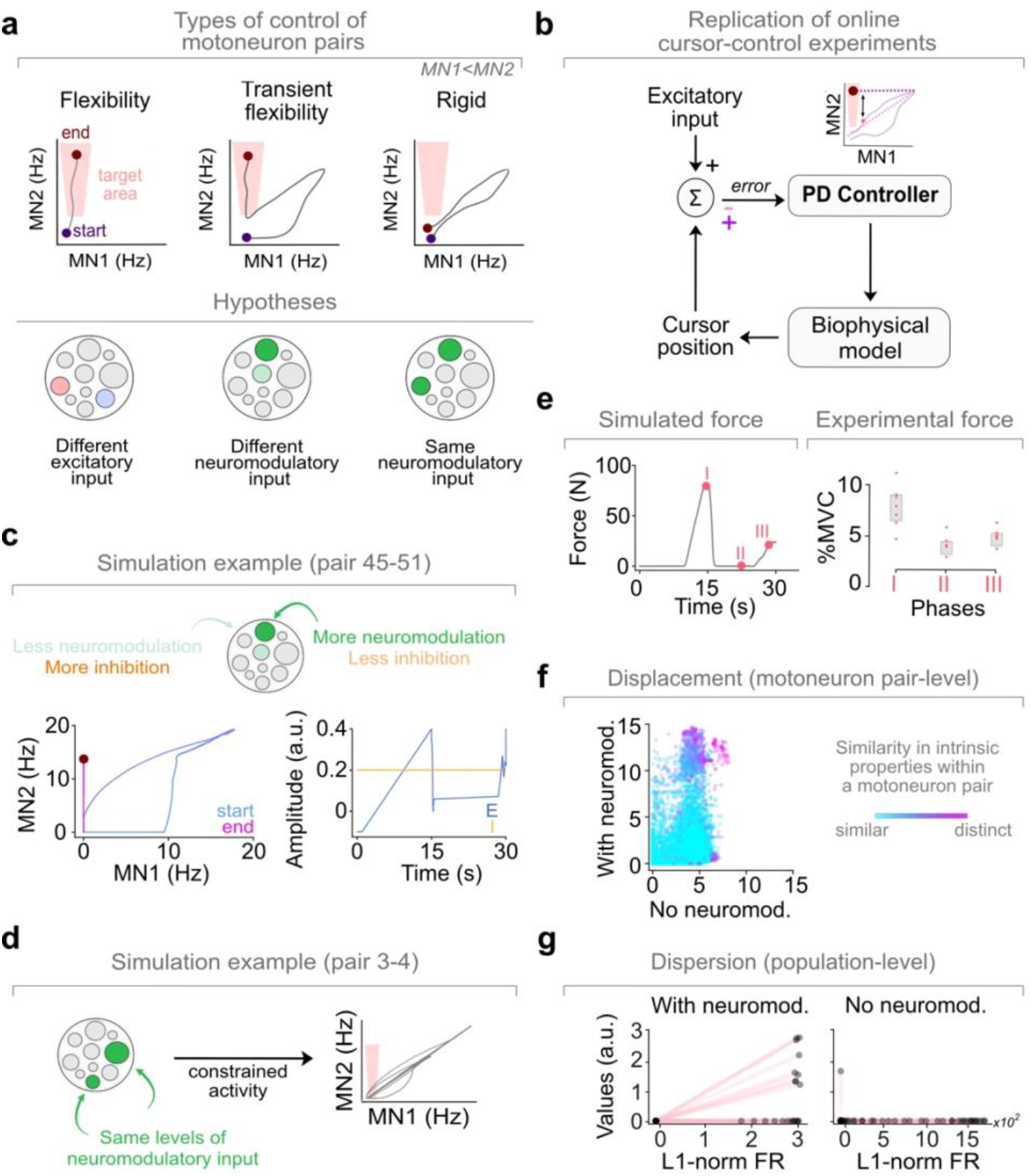
Simulations of the online cursor control experiments to decorrelate motoneuron pairs. **a:** In motoneuron control, full flexibility would imply independent control through different synaptic inputs, whereas rigid control would arise when these inputs are the same (or shared). The participants in Bräcklein et al.^11^ modulated the firing rates of two motoneurons to bring the cursor towards a target (here, only shown the 90° target) by exploiting transient decorrelation during derecruitment. **b:** To replicate these experiments, a PD controller adjusted the shared excitatory input by computing an error signal reflecting the distance to the target. **c:** Simulation example for motoneuron pair 45–51 (Δ*V*_*thCa*_ = 3) for Participant 1. The differential effect of neuromodulation on these two motoneurons led to transient decorrelation during derecruitment, as indicated by the cursor trajectory (left). Right, time-varying shared excitatory (blue) and inhibitory (orange) synaptic inputs. **d:** Simulated cursor trajectory for motoneuron pair 3–4 (Δ*V*_*thCa*_ = 0). Motoneurons were similarly affected by neuromodulation, and their activity remained largely correlated, as is characteristic of rigid control. **e:** Time-varying force trajectories generated by the model (left) recapitulated the characteristic end-of-trial increase observed in experimental data (right). **f:** Displacement captured a decrease in correlation between the pair of controlled motoneurons with increased neuromodulation, which was greater when motoneuron intrinsic properties were more different (colorbar). **g:** Dispersion showed a limited motoneuron pool-wide effect of neuromodulation during the last 1 s of successful 90º trials.

We reproduced this task by having our controller modulate the firing-rates of MN1 and MN2 by adjusting the shared excitatory synaptic input to the entire motoneuron pool, so the cursor intercepted the 90º target. By construction, like in the experiments, this would require using this unique shared input^1,2^ to decorrelate the activity of MN1 and MN2. The controller determined the ongoing shared synaptic input based on two Euclidean distance signals measuring the error between the firing rates of MN1 and MN2 that time point and the target coordinates in the axis they each controlled at (Fig. 4b).

Simulations showed that decorrelation between MN1 and MN2 only occurred when the two controlled motoneurons had sufficiently different intrinsic properties (Fig. 4c shows a simulation example with motoneuron pair 45-51 —the number indicates their size ranking within the pool—, for which Δ*V*_*thCa*_ = 3; Suppl. Fig. 4 shows additional examples for the remaining seven participant-specific models). In contrast, when the controlled motoneurons had similar properties, their activity remained correlated (Fig. 4a, “rigid control”), preventing the controller from acquiring the 90º target because the cursor trajectory remained confined to a straight line along the workspace (Fig. 4d, simulation example with motoneuron pair 3-4, where Δ*V*_*thCa*_ = 0). This result held across all three simulated relationships between excitatory and inhibitory synaptic inputs (Suppl. Fig. 4).

Having established that our model could leverage the effects of hysteresis to acquire the 90º target by exploiting the persistent firing of the larger motoneuron MN2, we tested the prediction that increasing the effect of neuromodulation should increase deviations from rigid motoneuron control. We computed two complementary metrics, displacement, and dispersion^5^. Displacement quantifies how far the firing rates of a pair of motoneurons deviates from perfectly correlated behavior. Dispersion quantifies how much the motoneuron pool firing pattern can vary while producing the same overall activity.

According to both metrics, increased neuromodulation led to greater transient flexibility, but this flexibility occurred only during derecruitment. Moreover, neuromodulation only produced larger pairwise displacement values —that is, greater decorrelation— when motoneurons differed in their intrinsic properties (Fig. 4f). However, dispersion values computed over the last second of the trial showed that the effect of neuromodulation at the population level remained confined to a subset of motoneurons and emerged only at the end of the trial (Fig. 4g). In contrast, dispersion values computed over the full trial remained close to zero both with and without neuromodulation (Suppl. Fig. 5). Thus, even when neuromodulation enabled transient decorrelation between pairs of motoneurons with different biophysical properties, the overall activity of the motoneuron pool remained largely unchanged.

These simulations were based on having our controller modulate the shared synaptic input to acquire the 90º target. But did the controller deploy the same strategy as participants of the original study did^11^? To verify that this was the case, we compared the simulated muscle force to the experimentally measured ankle force. Participants consistently acquired the 90º target by rapidly increasing their muscle force, and then decreasing it and increasing it again^11^ (Fig. 4e) —we refer to these three phases of force generation as phases I, II and III. Interestingly, our model naturally found this same behavioral strategy (Fig. 4e), suggesting that it is an effective way to achieve the task under the assumed physiological constraints. In summary, our model confirmed the prediction that neuromodulation enables the transient decorrelation of motoneuron pairs during derecruitment, even when a motoneuron pool shares the same excitatory synaptic input. Consistent with experimental observations, achieving this transient flexibility is only possible for pairs of motoneurons with sufficiently different intrinsic properties.

### Refined force control strategies can increase the transient flexible control of motoneurons enabled by neuromodulation

During the online cursor control experiments, participants improved their performance on the 90º target over 14 consecutive days of practice^11^, with their average success rate increasing from 33.3 ± 30.4% in the initial sessions to 65.0 ± 31.6% in the final sessions (Mann–Whitney U = 34.0, *P* = 0.03) (Fig. 5a, left). Here, we hypothesized that this behavioral improvement should result from a refined behavioral strategy whereby participants learned a better way to transiently decorrelate motoneuron activity, as opposed to reflect changes in the underlying physiological substrate (e.g., gradual adjustments in neuromodulatory input over days) which is unlikely to happen under these conditions.

**Fig. 5.**
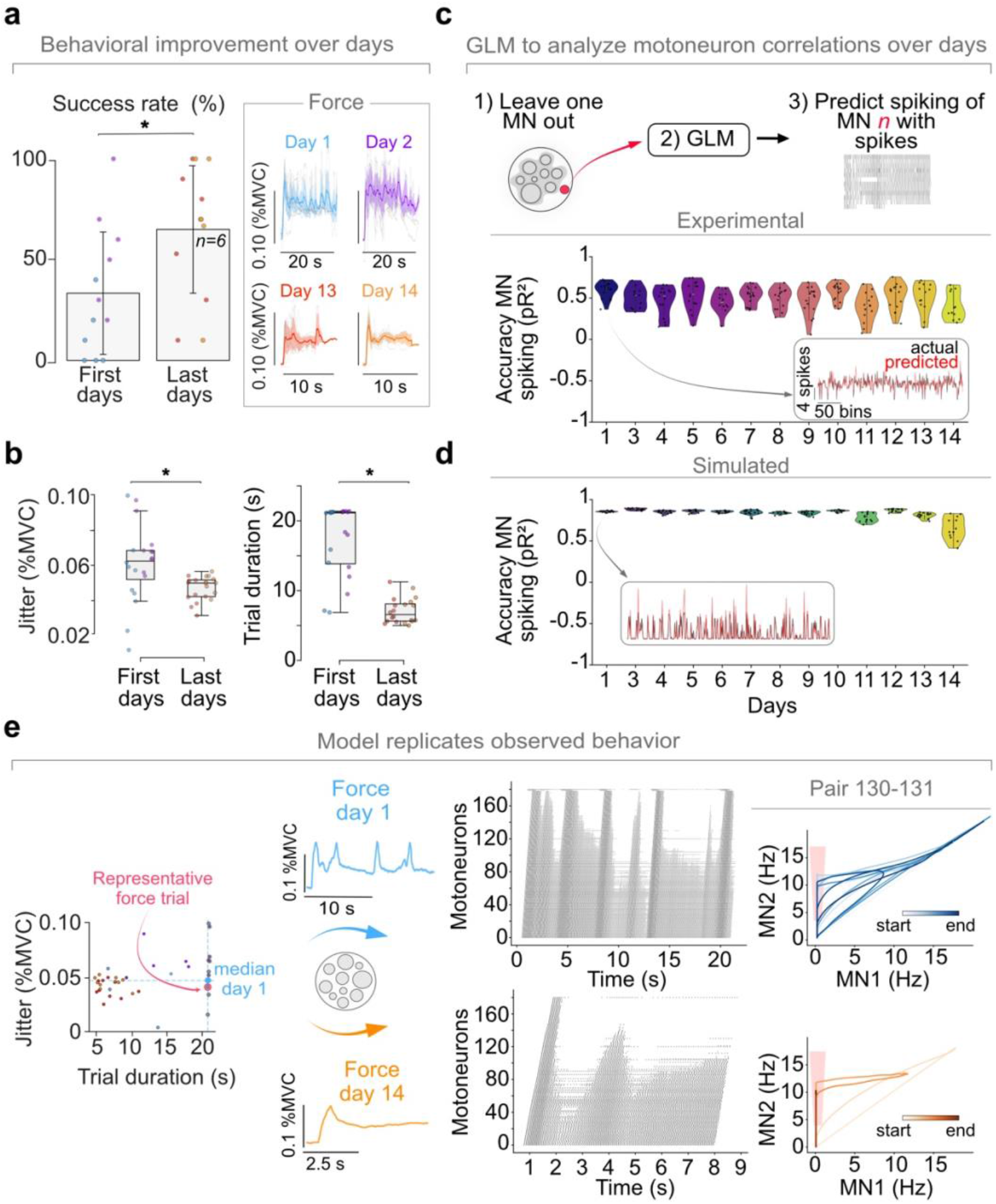
Improvements in flexible motoneuron control with practice arise from a refined behavioral strategy, rather than from physiological changes. **a:** Behavioral performance across training days. Left: success rates across participants (n = 6) for early (days 1–2) and late (days 13–14) sessions. Markers, one session from Participant 1. Right: average force trajectories (as % MVC) during the first and last two training days for a representative participant. Note the increased smoothness and greater consistency across trials. **b:** Decrease of force jitter (left) and trial duration (right) with practice. Data presented as in (a). **c:** Poisson GLM analysis to predict the spiking of each motoneuron from the spiking of the other recorded motoneurons. Experimental data show no systematic change in prediction quality across days. Inset, example predictions. **d:** Same as (c) but for simulated spiking activity using the recorded force as a proxy for shared synaptic input. **e:** Representative trials from the first (Day 1) and last (Day 14) session for Participant 1. Note the greater variability in motoneuron activity (rasters in the middle showing the activity of the entire pool) and cursor trajectory (right) early in learning. This example shows two motoneurons with different intrinsic properties: motoneurons 130– 131 (Δ*V*_*thCa*_ = 0.66). We show the most representative trial for each session, defined as the one closest to the median of the join force jitter – trial duration distribution for that session.

We first confirmed that improved performance was associated with clear changes in behavioral strategy by analyzing the original 14-day dataset^11^. Indeed, force profiles and cursor trajectories showed reduced variability during the late sessions as reflected by their reduced jitter (Mann–Whitney U = 314, *P* = 0.002), whereas trial duration decreased across days (Mann–Whitney U = 380, *P* <0.001), and the percentage of successful trials increased (Fig. 5a,b). Moreover, in the late sessions, trials not only became shorter and the cursor trajectories smoother, the force profiles became more stereotyped (Fig. 5a,b). These changes were consistent with improved control in all but two participants.

Having observed this refinement of force trajectories associated with improved task performance, we asked whether these behavioral changes could explain the increased decorrelation between motoneuron pairs during derecruitment that was associated with greater success rate. If behavioral changes drove the greater hysteresis in late experimental sessions, then the statistical relationships across motoneurons in the population should remain largely consistent across days. In contrast, if behavioral improvements were also associated with changes in the physiological substrate^35^ that enabled, e.g., more targeted effects of neuromodulation, these statistical relationships should weaken over time. We tested whether the statistical relationships between motoneurons changed across days using a Poisson Generalized Linear Model (GLM)^36–38^. For each motoneuron, the spiking activity during the entire trial was predicted from the spiking activity of the remaining motoneurons. As expected, model predictions measured by their Pseudo-R^2^ (Ref. 37,39) remained stable across days, with no clear downward trend (Fig. 5c; Suppl. Fig. 6a). This result also held when we used muscle force as input to predict the spiking of each motoneuron, under the assumption of a fixed relationship between the two (Suppl. Fig. 6b). Therefore, our GLM analyses indicate that participants’ improved ability to sustain the firing of the larger motoneuron during derecruitment is associated with a refined behavioral strategy, without requiring changes in motoneuron-level physiology.

To test more directly that an improved behavioral strategy alone could reproduce the increased motoneuron decorrelation across days, we simulated the behavior of the motoneuron pool given the experimentally measured force for every session of each of the six participants. We used the recorded force trajectories as a proxy of the time-varying shared excitatory synaptic input^40^ to the model, which then generated simulated time-varying motoneuron spiking. We first verified that the same GLM analysis used for the experimental data showed no systematic trend in prediction quality across days, observing that the predictions again remained accurate, ruling out large underlying physiological changes (Fig. 5d, Suppl. Fig. 6; Suppl. Fig. 7 confirms that the generated firing-rate distributions matched experimental values). Examining the simulated motoneuron activity patterns shows their similarity to the experimental data: during the early sessions, the model generated more variable spiking patterns (Fig. 5e, top; Fano factor, day 1 = 29.6 vs. day 14 = 21.8), and produced less directed cursor trajectories (Fig. 5e, top shows a pair of motoneurons, 130-131, that differed in their sensitivity to neuromodulation; Δ*V*_*thCa*_ = 0.66). In contrast, for the final days of practice the model reproduced the smoother and more directed cursor trajectories observed experimentally (Fig. 5e, bottom) without requiring any change in how neuromodulation affected the simulated motoneuron pool. Thus, improvements in task performance with practice reflect behavioral refinements rather than physiological changes affecting how neuromodulation affects motoneuron activity.

## Discussion

Our results show that the heterogeneous effects of neuromodulation on motoneurons from the same pool can enable transient flexibility during recruitment, despite strong physiological constrains that typically result in correlated firing. This flexibility arises through the differential effects of coupled neuromodulatory and inhibitory processes defining motoneuron activity together with a shared excitatory synaptic input. When challenged, people can learn to exploit this transient flexibility by refining their behavioral strategies over days of practice.

### Flexible control of motoneuron populations

Here, we have investigated the *flexibility* of motoneuron control under the most constrained regime: within a single motoneuron pool that receives a shared synaptic input^1,2^. This is different from recent studies^5,9^ that have focused on muscles that presumably receive multiple shared inputs, since the recruitment order of motoneuron pairs changed across conditions. This observation stands in stark contrast with the participants from Bräcklein et al.^11^, who failed to reverse the recruitment order of pair of motoneurons, even after many days of practice —with the caveat that these motoneurons were not the same across days. Remarkably, in monkeys, intracortical stimulation of different sites within primary motor cortex could reverse the recruitment order of motoneurons innervating the same proximal arm muscle^5^. Likewise, participants engaged in motoneuron control experiments like those in the leg that we have simulated here could selectively recruit different motoneurons from the biceps brachii^9^. Given this selective activation of specific motoneurons, it is likely that this flexibility arose upstream of the motoneuron pool through anatomical segregation of synaptic inputs. This may be due to the fact that the study that we have simulated involved leg muscles, and the experiments revealing flexible recruitment focused on the arm, which receives richer descending input from cortex^41,42^, although the relatively constrained nature of our task may have also contributed.

To directly test the flexibility of motoneuron control in the presence of a shared synaptic input, Bräcklein et al.^11^ asked participants to decorrelate two motoneurons with similar recruitment thresholds, a situation that, under the rigid control view, would make selective control particularly difficult (Fig. 4a). Participants could not selectively activate the higher-threshold (larger) motoneuron during recruitment, but consistently produced transient decorrelations during derecruitment (Fig. 5a). Our model suggests an explanation for these transient departures from rigid control: the heterogeneous effect of coupled neuromodulatory and inhibitory inputs alter motoneuron gain, producing transient decorrelations even in the presence of a shared excitatory synaptic input (compare Fig. 4c and Fig. 4d). Interestingly, neuromodulation led to the decorrelation of only a subset of motoneuron pairs (Fig. 4f), an effect that was largely washed out when looking at the entire pool (Fig. 4g, Suppl. Fig. 5). Therefore, neuromodulation seems to have a limited effect on how an entire motoneuron pool coordinates as a whole.

Even under these constraints, when explicitly challenged, participants could learn to increase the decorrelation between motoneuron pairs during derecruitment (Fig. 5a). We hypothesized that given the structure of neuromodulatory input to motoneurons^19^ and the “rigidity” of the descending pathways innervating motoneurons^43,44^, this improved performance would be achieved without changes in the underlying physiological substrate. We tested this hypothesis by building Poisson GLMs that predicted the activity of left out motoneurons from the rest of the population. If there were changes in the underlying physiological substrate, the performance of these models should degrade drastically, as directly shown by the lack of GLM generalisation following pharmacological manipulations of the crab stomatogastric circuit^35^. The consistent performance of our GLMs across days (Fig. 5c,d) allowed us to rule out a large change in the underlying substrate. Moreover, additional simulations in which we modelled the response of a fixed pool of motoneurons to generate force that participants produced over the different sessions further suggested that their refined behavioral strategies alone (force profiles; Fig. 5e) were sufficient to increase motoneuron decorrelation during derecruitment.

### Physiological interactions across motoneuron inputs

Our simulations show that only by having non-uniform sensitivity to neuromodulatory and inhibitory input we can reproduce the experimentally observed flexibility (Fig. 4d, Suppl. Fig. 4), also emphasizing the importance of inhibitory synaptic inputs. Because inhibition counteracts motoneuron excitability^13,23,24^, a uniform inhibitory input fails to reproduce the changes in motoneuron order during derecruitment observed in experiments (Fig. 4c). Yet, our results held across different simulated relationships between inhibition and neuromodulation^26^ (Suppl. Fig 4) all of which assumed a heterogeneous distribution of inhibitory input, consistent with recent experimental observations^17^. How inhibition and neuromodulation interact remains an open question, but their balance likely determines the physiological limits of motoneuron decorrelation during derecruitment. Future work could test this directly, for example, by transiently increasing inhibition via reciprocal inhibition of the tibialis anterior and assessing whether task performance declines.

### Limitations and future work

Our model captures key biophysical features of human motoneurons but simplifies several components of spinal circuitry. For instance, neuromodulatory effects were represented through calcium-mediated PICs, although other conductances, such as those capturing persistent sodium currents^45^, also contribute to persistent firing and were not explicitly modelled. We also did not include recurrent inhibition, Ia/Ib afferent pathways, or heteronymous pathways, all of which shape motoneuron excitability^16,46,47^ and thus can interact with neuromodulatory inputs in ways not represented here.

The simulated pool consisted mainly of slow-type motoneurons, reflecting the portion of the pool that is active at the low forces required for the online cursor-control task that we simulated^11^. Including fast-type, higher-threshold motoneurons could reveal additional interactions between neuromodulation, motoneuron recruitment dynamics, and flexible control. Finally, neuromodulatory and inhibitory inputs were implemented as static heterogeneous profiles, even if they are known to fluctuate with arousal, behavioral state, and interneuron activity^48^. The simplified dendritic morphology and the absence of dynamic inhibitory circuits may omit other factors that could also influence transient decorrelation. These limitations could be addressed in future experiments that directly manipulate inhibitory pathways or neuromodulatory input to causally test how inhibition and neuromodulation jointly set the physiological limits of flexible motoneuron control.

### Conclusion

In summary, our study shows that the heterogeneous effect of neuromodulation across motoneurons from the same pool is necessary and sufficient to account for their transient decorrelation during derecruitment. This transient flexibility emerges within an otherwise rigidly constrained framework, and can be better exploited with repeated practice. Besides shedding light on the effects of neuromodulation on motoneuron control, this work identifies fundamental challenges to develop motoneuron-based approaches for “human augmentation”^49^ that seek to leverage multiple control signals from the same rigidly controlled muscle.

## Methods

### Biophysical model

#### Architecture

We implemented a biophysical motoneuron model based on Hodgkin–Huxley equations, consisting of two electrically coupled compartments representing a soma and a dendrite^50^. The dendritic compartment included excitatory synaptic input, inhibitory synaptic input, leak conductance, and voltage-dependent calcium channels, which are critical for generating neuromodulatory effects^21,22^. Calcium channels were defined with a voltage deactivation threshold *V*_*thCa*_ distinct from the action potential threshold (*V*_*AP*_), enabling sustained firing (hysteresis)^14^. The somatic compartment included leak, fast potassium, slow potassium, and sodium channels. The soma and dendrite were coupled through an axial conductance, allowing symmetrical bidirectional current flow between compartments. The coupling conductance *g*_*c*_ was defined as:

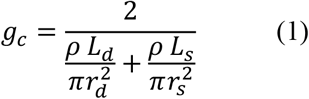

where *ρ* is intracellular resistivity, and *L*_*d*_, *L*_*s*_, *r*_*d*_, and *r*_*s*_ denote dendritic and somatic lengths and radii, respectively. Morphological and passive membrane parameters, including intracellular resistivity and compartment lengths and radii, are reported in Supplementary Table 1.

The somatic membrane potential (*V*_*s*_) evolved according to:

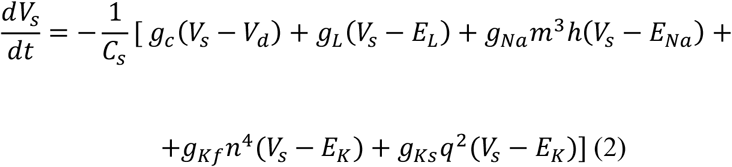

The dendritic membrane potential (*V*_*d*_) evolved according to:

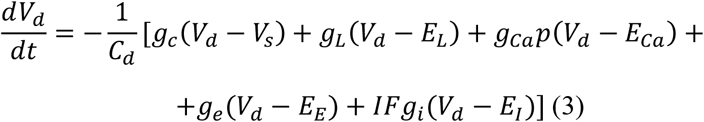

Here, *C*_*s*_ and *C*_*d*_ denote somatic and dendritic membrane capacitances. *g*_*L*_ denotes leak conductance; *g*_*Na*_, *g*_*Kf*_, and *g*_*Ks*_ denote sodium, fast potassium, and slow potassium conductances in the soma; and *g*_*Ca*_ denotes dendritic calcium conductance. *g*_*e*_ and *g*_*i*_ represent excitatory and inhibitory synaptic conductances, respectively. *E*_*L*_, *E*_*Na*_, *E*_*K*_, *E*_*Ca*_, *E*_*E*_, and *E*_*I*_ denote the corresponding reversal potentials. Inhibitory input was scaled by a multiplicative factor (*IF*), which was treated as an adjustable model parameter and optimized based on experimental data (see below). Specifically, the *IF* was defined to vary inversely with threshold *V*_*thCa*_, such that motoneurons with larger *V*_*thCa*_ values (stronger neuromodulatory effects) received proportionally less inhibition. After normalization across the pool, the largest *V*_*thCa*_ corresponded to an *IF* of 1, whereas the smallest *V*_*thCa*_ corresponded to an *IF* of 0.

All differential equations were solved using a fourth-order Runge–Kutta method with a fixed time step of 0.05 ms, corresponding to a sampling frequency of 20 kHz.

Gating variables *m, h, n, q*, and *p* followed first-order Hodgkin–Huxley kinetics, representing the probabilistic dynamics of ion channel activation and inactivation. Each gating variable evolved according to voltage-dependent forward and backward rate constants, *α*_*x*_(*V*) and *β*_*x*_(*V*), which governed channel opening and closing:

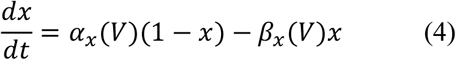

Where *x* ∈ {*m, h, n, q, p*}. Activating gates (*m, n, q*, and *p*) controlled channel opening, whereas the inactivating gate (*h*) governed sodium channel inactivation.

Action potential initiation occurred when the somatic membrane potential exceeded a firing threshold (*V*_*AP*_) defined as the steady-state depolarization produced by the rheobase current, given the motoneuron’s input resistance.

Neuromodulatory effects were implemented through dendritic calcium PICs^21,22^. Sustained firing was maintained when the somatic membrane potential remained above an effective threshold reduced by calcium channel activation:

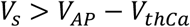

Where *V*_*thCa*_ *i*s a phenomenological parameter capturing the depolarizing contribution of calcium-mediated PICs. Incorporating *V*_*thCa*_ effectively lowered the firing threshold; larger values of this parameter produced stronger firing-rate hysteresis. The value of *V*_*thCa*_ was not fixed a priori and was optimized based on experimental data (see below). The calcium threshold *V*_*thCa*_ and the inhibitory scaling factor (*IF*) were not optimized independently but were constrained to vary inversely, reflecting the opposing effects of neuromodulation and inhibition on motoneuron excitability. Because inhibition counteracts motoneuron excitability^13,23,24^, a uniform inhibitory input was insufficient to reproduce the experimental data.

When motoneurons are repeatedly activated during successive contractions, their firing-rates increase, a phenomenon known as facilitation^14^. To capture this behavior, motoneurons that reached threshold during the ramp-down phase of a contraction were allowed to re-initiate firing once the above condition was satisfied. This mechanism introduced additional nonlinearities in the motoneuron pool transfer function.

#### Data to build the model

We used experimental data from decerebrated cat preparations^51–55^, focusing on slow-type motoneurons, as our simulations targeted submaximal contraction regimes.

**Table 1.**
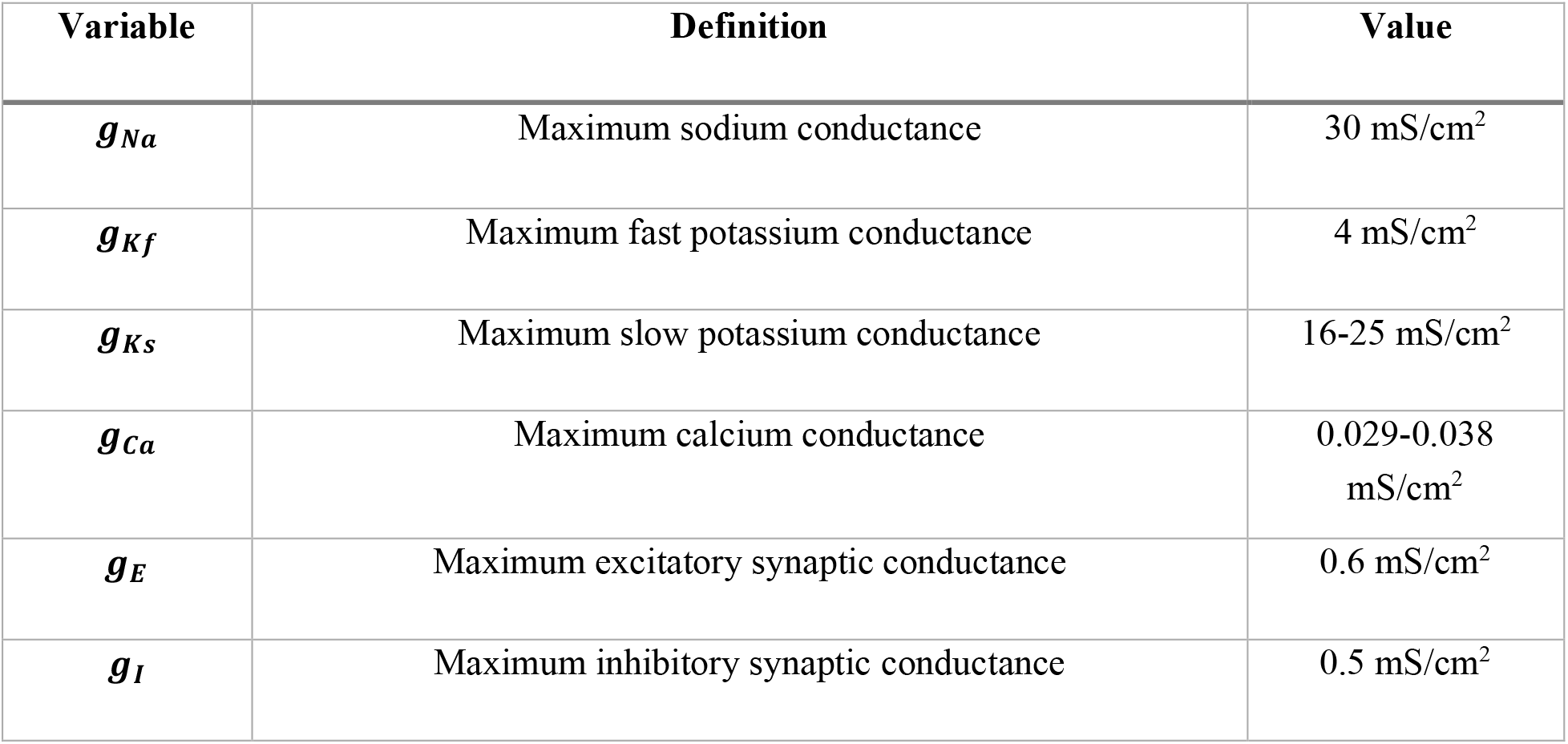
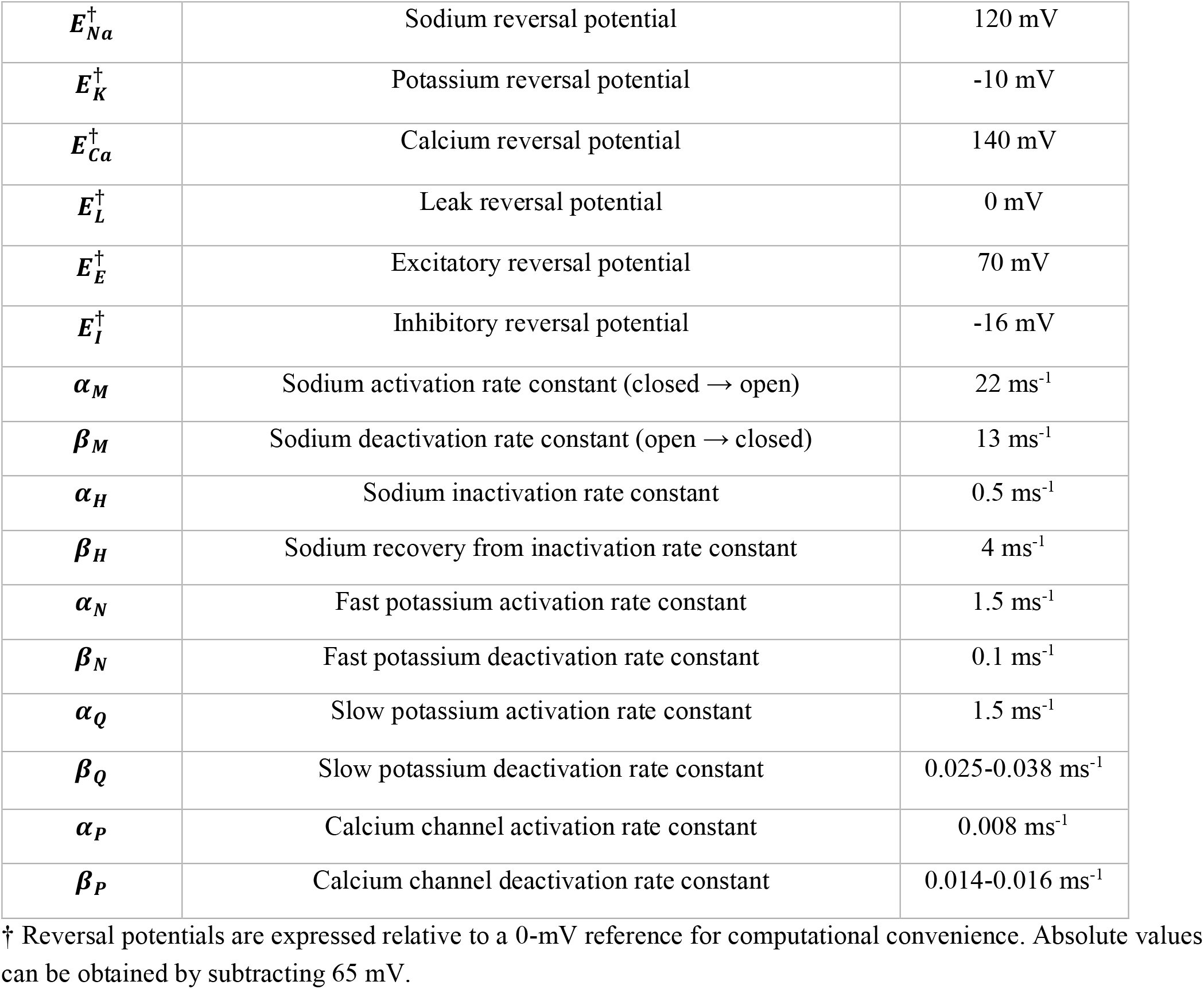
Core biophysical model parameters.

| Variable | Definition | Value |
| --- | --- | --- |
| $g_{Na}$ | Maximum sodium conductance | 30 mS/cm<sup>2</sup> |
| $g_{Kf}$ | Maximum fast potassium conductance | 4 mS/cm<sup>2</sup> |
| $g_{Ks}$ | Maximum slow potassium conductance | 16-25 mS/cm<sup>2</sup> |
| $g_{Ca}$ | Maximum calcium conductance | 0.029-0.038 mS/cm<sup>2</sup> |
| $g_E$ | Maximum excitatory synaptic conductance | 0.6 mS/cm<sup>2</sup> |
| $g_I$ | Maximum inhibitory synaptic conductance | 0.5 mS/cm<sup>2</sup> |
| $E_{Na}^{\dagger}$ | Sodium reversal potential | 120 mV |
| $E_K^{\dagger}$ | Potassium reversal potential | -10 mV |
| $E_{Ca}^{\dagger}$ | Calcium reversal potential | 140 mV |
| $E_L^{\dagger}$ | Leak reversal potential | 0 mV |
| $E_E^{\dagger}$ | Excitatory reversal potential | 70 mV |
| $E_I^{\dagger}$ | Inhibitory reversal potential | -16 mV |
| $\alpha_M$ | Sodium activation rate constant (closed → open) | 22 ms<sup>-1</sup> |
| $\beta_M$ | Sodium deactivation rate constant (open → closed) | 13 ms<sup>-1</sup> |
| $\alpha_H$ | Sodium inactivation rate constant | 0.5 ms<sup>-1</sup> |
| $\beta_H$ | Sodium recovery from inactivation rate constant | 4 ms<sup>-1</sup> |
| $\alpha_N$ | Fast potassium activation rate constant | 1.5 ms<sup>-1</sup> |
| $\beta_N$ | Fast potassium deactivation rate constant | 0.1 ms<sup>-1</sup> |
| $\alpha_Q$ | Slow potassium activation rate constant | 1.5 ms<sup>-1</sup> |
| $\beta_Q$ | Slow potassium deactivation rate constant | 0.025-0.038 ms<sup>-1</sup> |
| $\alpha_P$ | Calcium channel activation rate constant | 0.008 ms<sup>-1</sup> |
| $\beta_P$ | Calcium channel deactivation rate constant | 0.014-0.016 ms<sup>-1</sup> |

† Reversal potentials are expressed relative to a 0-mV reference for computational convenience. Absolute values can be obtained by subtracting 65 mV.

### Computation of hysteresis

Hysteresis refers to the sustained firing that persists after the cessation of synaptic input. It was quantified as the ratio between the force at derecruitment and the force at recruitment, normalized by subtracting one. Negative values indicated the presence of hysteresis. We did not use alternative measures such as delta-F^56^ because the motoneuron pool received a share synaptic input^11^, eliminating the need of a reference motoneuron to calculate the prolonged firing. This metric does not capture the absolute level of neuromodulation, but rather quantifies the prolonged firing activity resulting from neuromodulatory effects.

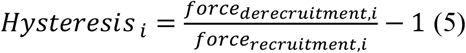

### Experimental data used for model optimization

The following text is a summary of the experimental protocol described by Avrillon et al^18^. Eight healthy participants (27 ± 3 years) performed isometric ankle dorsiflexions with the dominant foot secured to a custom ankle dynamometer. Participants were seated with the hip flexed at 45°, the knee fully extended, and the ankle fixed at 30° plantarflexion.

Tibialis anterior muscle activity was recorded using HDsEMG with four 64-channel electrode grids (256 channels in total) sampled at 2048 Hz. After completing maximal voluntary contractions (MVCs) for force normalization, participants performed eight submaximal trapezoidal contractions ranging from 10 % to 80 % MVC, presented in random order. Each contraction included ramp-up and ramp-down phases at a constant rate of 5 % MVC·s^−1^, with plateau durations of 10–20 s depending on the target force level.

Raw signals were band-pass filtered between 20 and 500 Hz using a second-order Butterworth filter to remove motion artefacts and high-frequency noise. Channels with poor signal quality (identified by excessive noise or low signal-to-noise ratio) were visually inspected and excluded from analysis.

Motoneuron spike trains were extracted using convolutive blind source separation^57^. Discharge times were automatically detected from the separated sources using peak detection and clustering and subsequently refined by manual inspection to remove artefacts or missed discharges. On average, 42 ± 24 motor units were identified per contraction, yielding 129 ± 44 unique motoneurons per participant.

Motoneurons were tracked across force levels based on the spatial distribution of their action potentials on the electrode array. Tracking was performed by comparing the root-mean-square error (RMSE) between average action-potential waveforms across conditions. Motoneurons were retained if their across-trial RMSE was lower than the 5^th^ percentile of the RMSE distribution computed with all other motoneurons, ensuring waveform uniqueness.

### Black-box optimization of calcium channel voltage deactivation thresholds

We employed a Black-Box evolutionary optimizer^27^ functioning as a hyperparameter sweep to estimate the *V*_*thCa*_ values responsible for producing different levels of hysteresis. The objective function minimized the sum of squared distances (SSD) between experimental and simulated hysteresis values:

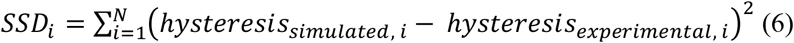

The optimization began by randomly initializing a population of *N* candidate solutions, each representing a possible *V*_*thCa*_ value within a predefined search range. This range was constrained through prior exploration using the Weights & Biases tool^58^, and we found that *V*_*thCa*_ values between 0 and 7 produced physiologically plausible behavior consistent with the size principle.

At each optimization run, new candidate solutions were generated by introducing mutant variants of existing *V*_*thCa*_ values, following a differential evolution strategy. Candidate solutions were evaluated based on their ability to minimize the SSD, and the best-performing solutions were retained.

For each motoneuron, the optimization was performed using a time-limited evolutionary search (maximum runtime of 200s per run). For motoneurons exhibiting hysteresis, the optimizer was restarted until convergence (SSD < 0.005). Across successive restarts, population size, mutation rate, and crossover rate were progressively increased to promote broader exploration of the parameter space and to reduce sensitivity to local minima (PopulationSize = 100 + 10*r*, MutationRate = 0.8 + 0.05*r*, CrossoverRate = 0.8 + 0.05*r*, where *r* denotes the restart index).

To quantify model fit, we computed the coefficient of determination (*R*^2^). The total sum of squares (TSS) was calculated from the average experimental hysteresis to measure overall variability, while the residual sum of squares (RSS) quantified the mismatch between experimental and simulated values. *R*^2^ was then computed as 1 − RSS/TSS. A separate model was fitted for each participant (*n* = 8) to capture individual variability in motoneuron derecruitment dynamics.

### Quantification of firing rate distribution overlap

Kernel density estimates (KDEs) were computed for each contraction level using a Gaussian kernel (SciPy gaussian_kde). All firing-rate values were pooled to define a common evaluation range, and each KDE was sampled over 500 uniformly spaced points. To assess the range in which firing-rate distributions overlapped across contraction levels, each KDE was normalized to its own maximum, and we evaluated whether it contributed meaningfully at each point on the shared grid. A distribution was considered active when its density exceeded 1% of its peak value (threshold chosen to suppress numerical noise in low-density regions). The overlap intensity was defined as the number of distributions active at each grid point, providing a non-parametric estimate of the firing-rate region jointly supported across contraction levels.

### Computation of firing rates

Motoneuron firing-rates were computed using two complementary approaches depending on the analysis. For precise individual estimates, firing-rates were derived from inter-spike intervals, defined as the time difference between consecutive discharges. For analyses requiring firing-rate vectors of equal length (such as state-space comparisons between motoneuron pairs), binary spike trains were convolved with a 400-ms Hanning window. This procedure smoothed discrete spike events into continuous firing-rate profiles while preserving the temporal structure of the discharges. The 400-ms window provided an optimal balance between temporal resolution and noise reduction^59,60^.

### Computation of force

The twitch model was implemented as a second-order critically damped system representing the discrete-time impulse response^20,61^. This formulation effectively captured the rapid rise and smooth decay of motoneuron responses, characteristic of twitch contractions in muscle fibers. The generated force *F*(*n*) was determined by the twitch amplitude (*A*_peak_), contraction time (*t*_peak_), time step (*T*), and the spike train (*E*(*n*)). Force was computed for each motoneuron individually (Eq. 7), and then summed across the pool to obtain the total muscle force.

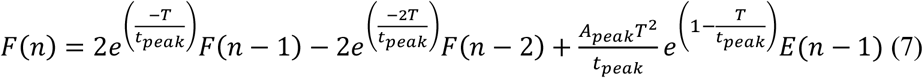

For each motoneuron, twitch parameters were sampled from predefined ranges to capture physiological variability: *A*_*peak*_ from 0.103–0.123, *t*_*peak*_ from 100–110 ms, and tetanic force limits from 0.393–0.491 (a.u.)^51–55^.

### Controller to replicate for tracking task

To assess the model’s ability to replicate force-tracking tasks, we implemented a PD controller. The controller used proportional and derivative gains of *k*_*p*_ = 2and *k*_*d*_ = 0.1, with a fixed update interval of Δ*t* = 0.05s. These values were selected through a parameter sweep to optimize tracking accuracy while minimizing oscillations in the controller response.

At each time step *i*, the tracking error was defined as the Euclidean distance between the target and simulated force trajectories:

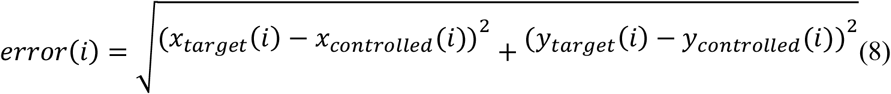

The control output was computed as:

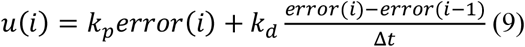

The controller output was scaled before being applied to the muscle model to keep excitation within physiological bounds. The controller was iterated for 25 passes to minimize residual tracking error while avoiding oscillations. The adjusted excitatory input was subsequently transformed into force output using the twitch filter and tetanic limits described above.

### Metrics for assessing flexibility in motoneurons

We implemented two metrics to evaluate flexibility of motoneurons, displacement and dispersion, as defined by Marshall et al.^5^ Under rigid control, motoneuron firing-rates are expected to evolve monotonically with a common input, such that increases in one motoneuron are accompanied by non-decreasing activity in all others. Deviations from this monotonic structure indicate flexible control.

**Displacement** measures the minimal adjustment required to make changes in population firing-rates consistent with monotonicity. For firing-rate vectors at two time points *t* and *t*′:

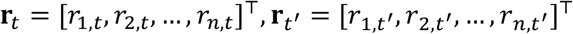

Displacement is defined as:

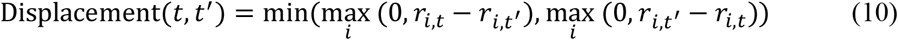

Displacement was computed over relative time lags *τ, τ*′ ∈ [−25,25]ms to account for differences in motoneuron response latencies, and the minimum value across all lagged comparisons was retained. Low displacement values indicate near-monotonic population dynamics, whereas higher values reflect stronger violations of rigid control.

**Dispersion** quantifies variability in population firing patterns at a fixed level of total activity. Total population activity was defined by the *L*_1_-norm ∥ **r**_*t*_ ∥_1_= ∑_*i*_ ∣ *r*_*i,t*_ ∣. Under rigid control, a given total activity level *λ* should correspond to a unique population firing pattern. Dispersion at activity level *λ* was therefore defined as the maximum *L*_1_-distance between any two population states with total activity within a small tolerance *ε* of *λ*, allowing for small temporal shifts *τ*_1_, *τ*_2_:

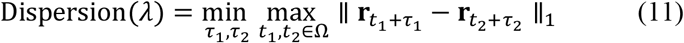

Where Ω = {*t*: ∥ **r**_*t*_ ∥_1_≈ *λ*}. A dispersion of zero indicates identical firing patterns at that activity level, consistent with perfectly rigid control.

### Data from online cursor-control experiments

We analyzed experimental data from the protocol described by Bräcklein et al.^11^. Eight healthy participants were originally recruited to learn volitional control of individual motoneurons in the tibialis anterior muscle, but in the present study we only included six.

Because our analyses required a sufficiently large and consistently recruited motoneuron population, both for assessing population-level effects and for fitting Poisson GLMs, these two participants were excluded from further analysis. Two participants exhibited atypical trends: Participant 6 showed minimal improvement across days, with success rates remaining low (~10% early vs. ~15% late sessions), whereas Participant 5 achieved higher success rates (35% early vs. 75% late sessions) despite increasingly noisy force traces and shorter trial durations, suggesting a distinct control strategy.

Participants were seated in a reclined position with their dominant foot secured to a custom ankle dynamometer (hip flexed 45°, knee fully extended, ankle fixed at 30° plantarflexion) to restrict overt movements and isolate tibialis anterior activation. HDsEMG was recorded from the tibialis anterior muscle using a 64-channel electrode grid, sampled at 12.2 kHz. The signals were decomposed online with a convolutional blind source separation algorithm, providing real-time discharge timings of individual motoneurons. After an initial calibration phase in which motoneurons were automatically identified and manually refined, a subset of well-isolated motoneurons was selected for neurofeedback control.

Participants then performed a two-dimensional online control task in which the firing-rates of two selected motoneurons (e.g., one lower-threshold and one higher-threshold) were mapped to orthogonal cursor axes. They were instructed to reach three types of visual targets: (i) activating only the lower-threshold motoneuron, (ii) activating both motoneurons together, or (iii) selectively recruiting the higher-threshold motoneuron. Firing-rates were estimated in real time with 800 ms sliding windows and normalized to the maximum rate observed during calibration. All contractions were performed at approximately 10% of MVC, ensuring that control relied on fine modulation rather than large force production. In the present study, we focused on the third target condition, as it provides the most direct test of flexible motoneuron control.

### Simulations of online control experiments

To replicate the experimental task described in Bräcklein et al.^11^, we implemented a PD controller that operated on pairs of motoneurons with similar recruitment thresholds (defined as motoneurons within five positions in the recruitment order). In the simulations shown, we focused on two representative motoneurons that differed in their levels of neuromodulatory input, as determined by their respective *V*_thCa_ values. For the example in the main text, the controller used proportional and derivative gains of *k*_*p*_ = 4 and *k*_*d*_ = 1, updated at a timestep of Δt = 0.1. For each participant, however, the gains were individually tuned by sweeping across candidate parameter combinations and selecting those that maximized decorrelation. The values used in each case are reported below.

For Participants 2–8, we analyzed one motoneuron pair per participant, with one motoneuron receiving stronger neuromodulatory input and weaker inhibition than the other. The gains used for each participant and condition were as follows (timestep Δt = 0.1 throughout). For Participant 2, the inversely proportional condition used *k*_*d*_ = 0.5 and *k*_*p*_ = 0.2, while both the proportional and constant conditions used *k*_*d*_ = 1 and *k*_*p*_ = 4. For Participant 3, the inversely proportional condition used *k*_*d*_ = 0.5 and *k*_*p*_ = 0.2, the proportional condition used k_d = 1 and *k*_*p*_ = 3, and the constant condition used *k*_*d*_ = 1 and *k*_*p*_ = 4. For Participant 4, the inversely proportional condition used *k*_*d*_ = 0.5 and *k*_*p*_ = 0.2, the proportional condition used *k*_*d*_ = 1 and *k*_*p*_ = 3, and the constant condition used *k*_*d*_ = 1 and *k*_*d*_ = 4. For Participants 5, 6, 7, and 8, all three conditions used *k*_*d*_ = 1 and *k*_*p*_ = 4.

The excitatory input profile consisted of a triangular contraction that began at a negative value (–0.1) to ensure that motoneurons remained subthreshold at rest and were only recruited as the excitatory input increased. This common excitatory input was iteratively updated by the controller over 20 iterations, where each iteration corresponded to one full pass of the triangular waveform. Inhibition was not controlled by the PD loop; instead, inhibitory inputs were derived by applying proportional, inversely proportional, or constant couplings to the excitatory input.

Spike trains were generated for the two motoneurons under these inputs and converted into firing-rates. For computational efficiency, firing-rates were averaged over non-overlapping windows of 20 samples. The resulting two-dimensional trajectory (defined by the firing-rates of the motoneuron pair) was then compared with the task-defined target trajectories.

Two control objectives were implemented sequentially, mirroring strategies observed in human experiments. First, once both motoneurons were recruited, the controller minimized error relative to the diagonal trajectory, corresponding to synchronous co-activation and deactivation of the pair. The Euclidean error between simulated and target trajectories was computed as in Eq. 8, and the control output *u*(*i*) as in Eq. 9. During this diagonal phase, control outputs were scaled (×0.002) and subtracted from the common excitatory input (*excitation*_*i*_ = *excitation*_*i*_ − *u*_*i*_), thereby driving synchronous deactivation of both motoneurons.

Second, once the firing-rate of the lower-threshold motoneuron fell below 5 Hz, the controller switched to a vertical-trajectory objective. Here, the error was computed relative to the maximal firing-rate of the higher-threshold motoneuron, encouraging selective activation of this unit while maintaining the lower-threshold unit silent. In this phase, control outputs were scaled (×0.0007) and added to the excitatory input (*excitation*_*i*_ = *excitation*_*i*_ + *u*_*i*_), selectively increasing the firing of the higher-threshold motoneuron.

This design reproduced the two-stage control strategy observed experimentally: participants first reduced the firing of both motoneurons along the diagonal, and then selectively increased the firing of the higher-threshold motoneuron without reactivating the lower-threshold motoneuron.

### Metrics to assess performance gains with practice

Force-control performance was quantified using two complementary metrics.

#### Force jitter

It was defined as the proportion of high-frequency content in the force trace. For each trial, the raw force signal was transformed using a one-sided Fast Fourier Transform, and the amplitude spectrum was normalized by signal length. The smoothness metric was then computed as the sum of amplitudes above 5 Hz, a cutoff chosen to exceed the typical bandwidth of voluntary force production in humans^40,62,63^. Higher values indicated noisier and less controlled force output.

#### Trial duration

Trial duration was defined as the time required to successfully complete the task. Analyses were restricted to trials in which participants were required to reach targets demanding a decorrelation between motoneuron pairs. Longer trial durations were interpreted as reflecting poorer control performance.

### Poisson generalized linear model to assess motoneuron correlation across days

#### Model

To quantify correlations between motoneurons across days, we employed Poisson generalized linear models (GLMs)^36–38^. The models predicted the spike-count activity of a given motoneuron from two sets of predictors: (i) the spiking activity of the remaining motoneuron population, and (ii) external behavioral covariates, including the recorded force^39^. GLMs extend ordinary linear regression by assuming that the response variable follows a distribution from the exponential family. For neural data, a Poisson distribution is appropriate because spike counts are discrete and non-negative. This makes the Poisson GLM well suited for motoneuron spiking activity. For each time bin, the GLM computes a weighted linear combination of input features:

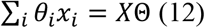

Where *X* contains the different covariates 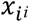, and Θ is the corresponding weight matrix for each of the covariates *θ*_*i*_.

This linear predictor is then passed through an exponential nonlinearity (inverse link function) to ensure the predicted firing-rate remains non-negative:

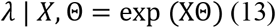

The observed spike count *n* in each bin is assumed to follow a Poisson distribution with mean *λ* ⋅ Δt where Δt is the bin duration:

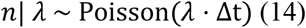

#### GLM Fitting

Using both the experimental and simulated data, we evaluated the ability of Poisson GLMs to predict the spiking activity of individual motoneurons based on either (1) the spiking activity of the remaining motoneurons or (2) force signals. Within each session, trials were concatenated across time to form a population matrix of size (motoneurons × time bins). Motoneurons with fewer than 10 spikes over the entire session were excluded to ensure that model fitting was performed only on sufficiently active motoneurons.

The response variable was defined as the spike count of a target motoneuron within each time bin. For predictors, either the activity of all other motoneurons (excluding the target) or the corresponding force values were used. To mitigate the influence of rare, abnormally large spike counts, bins exceeding the 99^th^ percentile of the target motoneuron’s spike distribution were excluded.

A bin width of 500 ms was used for all analyses. This value was determined empirically by testing a range of candidate widths and assessing how well the resulting spike count distributions conformed to a Poisson model. Goodness-of-fit was evaluated with chi-square tests^64^, where a significant result (*P* < 0.05) would indicate a deviation from Poisson statistics. A bin size of 500 ms yielded the most consistent agreement between observed and expected count distributions (e.g., χ^2^ = 2.99, *P* = 0.39).

Model fitting was performed using a fixed regularization strength (α = 0.1), chosen to balance bias and variance across datasets. To assess generalization, we employed 10-fold cross-validation. Folds with fewer than 10 bins or fewer than 5 non-zero spike counts were discarded to avoid unstable model estimation.

All GLMs were fit using the Newton–Cholesky solver, which is well-suited for regularised Poisson regression problems and provides a balance between computational efficiency and numerical stability^65^.

#### Evaluating performance

Model performance was quantified using a Poisson deviance-based pseudo-R^2^ (*pR*^2^) a relative measure of goodness-of-fit analogous to the coefficient of determination used in Gaussian models^37,39^. This metric compares the log-likelihood of the fitted model against a null model that predicts a constant firing-rate equal to the mean of the data.

The pseudo-R^2^ is defined as:

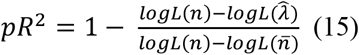

where *n* are the observed spike counts, 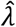 the model predictions and 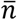 the mean of the spike count (the null model prediction). The Poisson likelihood of a spike train over *T* time bins is:

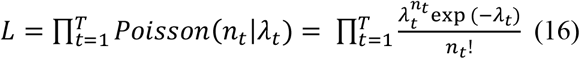

and the corresponding log-likelihood is:

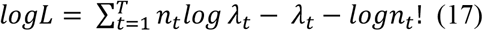

Then, a *pR*^2^ value of 1 indicates perfect prediction, whereas a value of 0 indicates no improvement over the null model. Negative values imply that the model performs worse than predicting the mean spike count.

### Statistics

Trial durations and force jitter were compared between early (first two days) and late (last two days) sessions using two-sided Mann-Whitney U tests. This non-parametric test was chosen as it does not assume normality of the data. Effect sizes were calculated as rank-biserial correlations, where values closer to ±1 indicate stronger group separation. Statistical significance was set at α = 0.05.

To assess changes in task success across training, success rates from early (first two days) and late (last two days) sessions were compared across all subjects using a two-sided Mann-Whitney U test. This non-parametric test was chosen as it does not assume normality of the data.

## Code availability

All the analysis will be made public upon publication in a peer reviewed journal.

## Data availability

Data will be made available upon reasonable request to the corresponding authors. Data from Avrillon et al. 2024 is public in https://doi.org/10.6084/m9.figshare.24640944.v1.

## Acknowledgements

This work was supported by the Imperial-META Wearable Neural Interfaces Research Centre (D.F. and J.A.G.), grant no. EP/T020970/1 from the UKRI Engineering and Physical Sciences Research Council (D.F. and J.A.G.), and grant no. ERC-2020-StG-949660 from the European Research Council (J.A.G.).

## Supplementary figures

**Supplementary Fig. 1.**
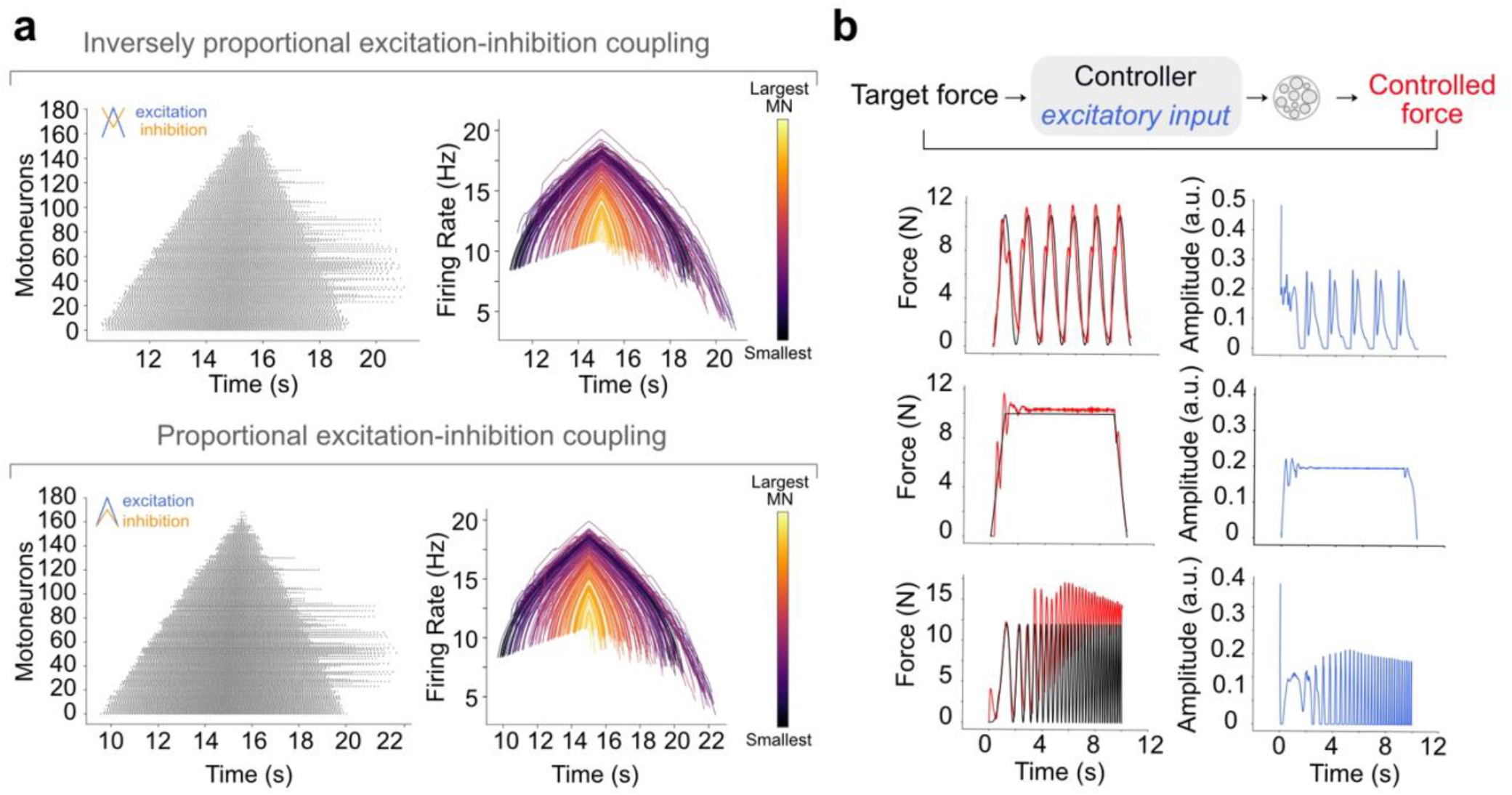
a: Different relationships between excitatory and inhibitory synaptic inputs recapitulate experimental motoneuron recruitment. Right: Inversely proportional relationship between inhibitory and excitatory synaptic inputs. Example raster plot and motoneuron population firing rates of one representative participant show orderly recruitment and variable derecruitment, including hysteresis. Left: Proportional relationship between inhibitory and excitatory synaptic inputs. Data presented as in (a). MN, motoneuron. **b: PD controller replicating force-tracking tasks**. The controller modulates the shared excitatory synaptic input to the motoneuron pool to minimize the instantaneous error between the simulated force and the target force (left). The controller reproduced the force output observed in humans (middle) across three tracking tasks, by generating the appropriate synaptic input to the motoneuron pool (right). These included, from top to bottom: (1) tracking a sinusoidal profile, (2) tracking a trapezoidal profile, and (3) attempting to track a high-frequency chirp signal, which the model eventually failed to do as it expected by the limited bandwidth of the neuromuscular system.

**Supplementary Fig. 2.**
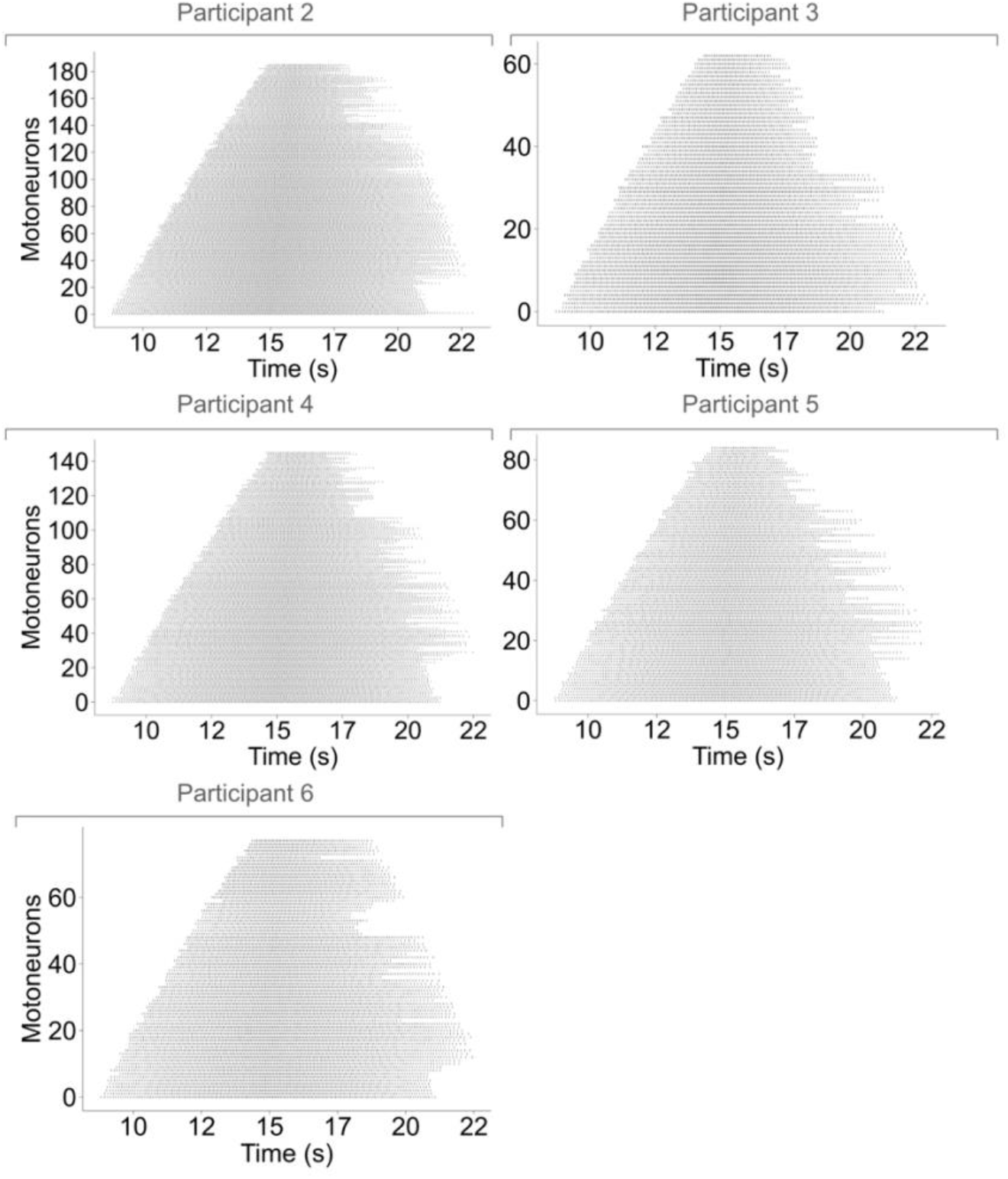
Simulated motoneuron activity for five additional participants. Raster plots consistently show orderly recruitment and variable derecruitment, including hysteresis.

**Supplementary Fig. 3.**
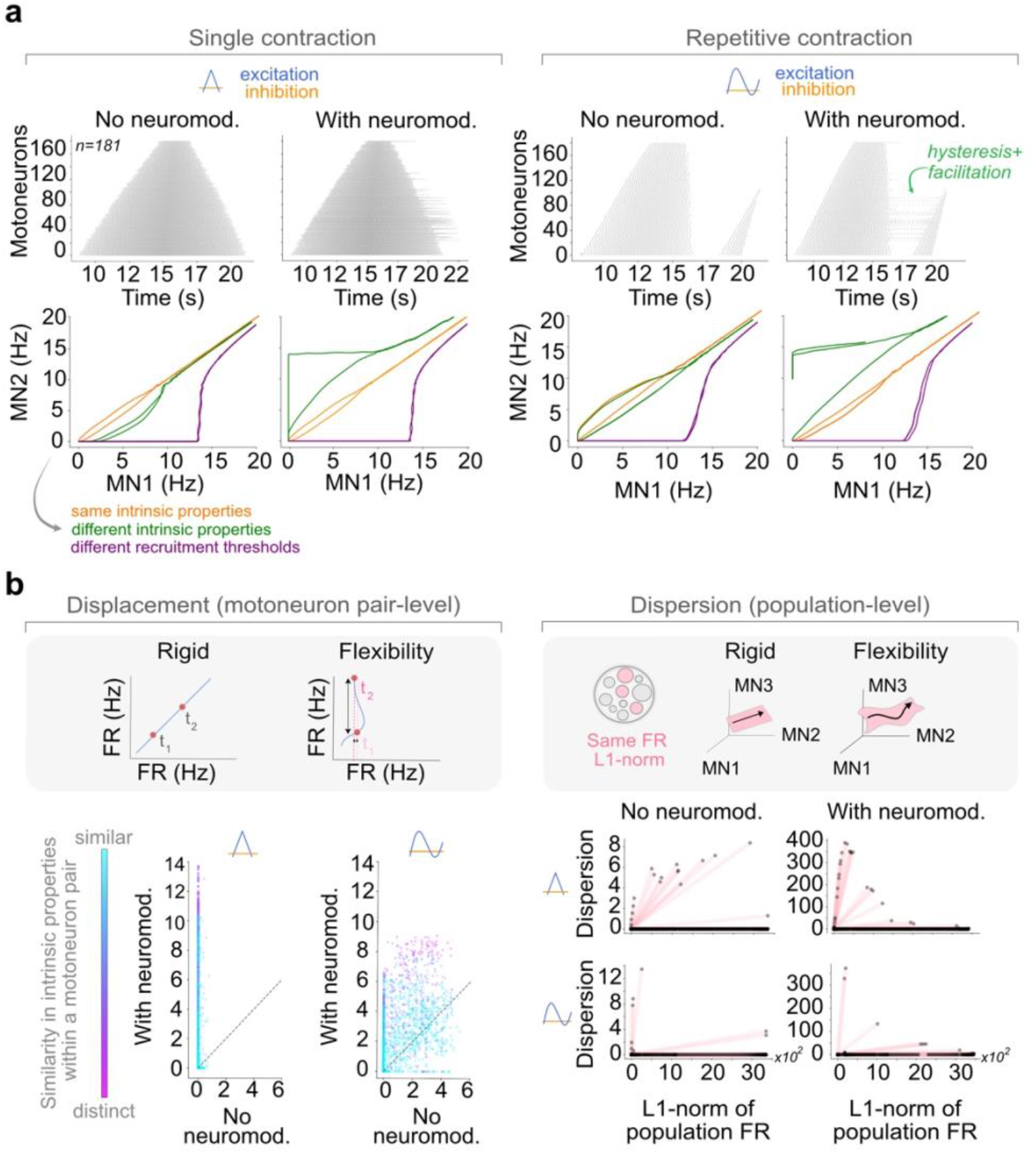
The effect of neuromodulation on the correlation between motoneuron pairs depends on the difference between their intrinsic properties, and always has little impact on the overall structure of population activity. **a:** Top, raster plots representing motoneuron activity during a single triangular contraction (left columns) and a dynamic sinusoidal contraction (right column). For both contractions, neuromodulation increased motoneuron variability and led to persistent firing (left-hand vs. right-hand side rasters). Bottom, state space trajectories of different motoneuron pairs reveal different behaviors that depend on how similar their intrinsic properties are. Only when a motoneuron pair had different intrinsic properties (V_thCa_), do their trajectories state-space deviated from the diagonal, reflecting distinct neuromodulatory inputs and reduced correlation. **b:** Flexibility metrics: displacement (deviation from correlated activity for a motoneuron pair), and dispersion (a measure of population-level flexibility) were computed for both contraction types. Neuromodulation and pairwise dissimilarity consistently increased displacement, with greater variability during dynamic contractions. Dispersion also increased with neuromodulation, but largely remained near zero. Thus, neuromodulation primarily enhances pairwise flexibility while exerting little influence on population-level dynamics.

**Supplementary Fig. 4.**
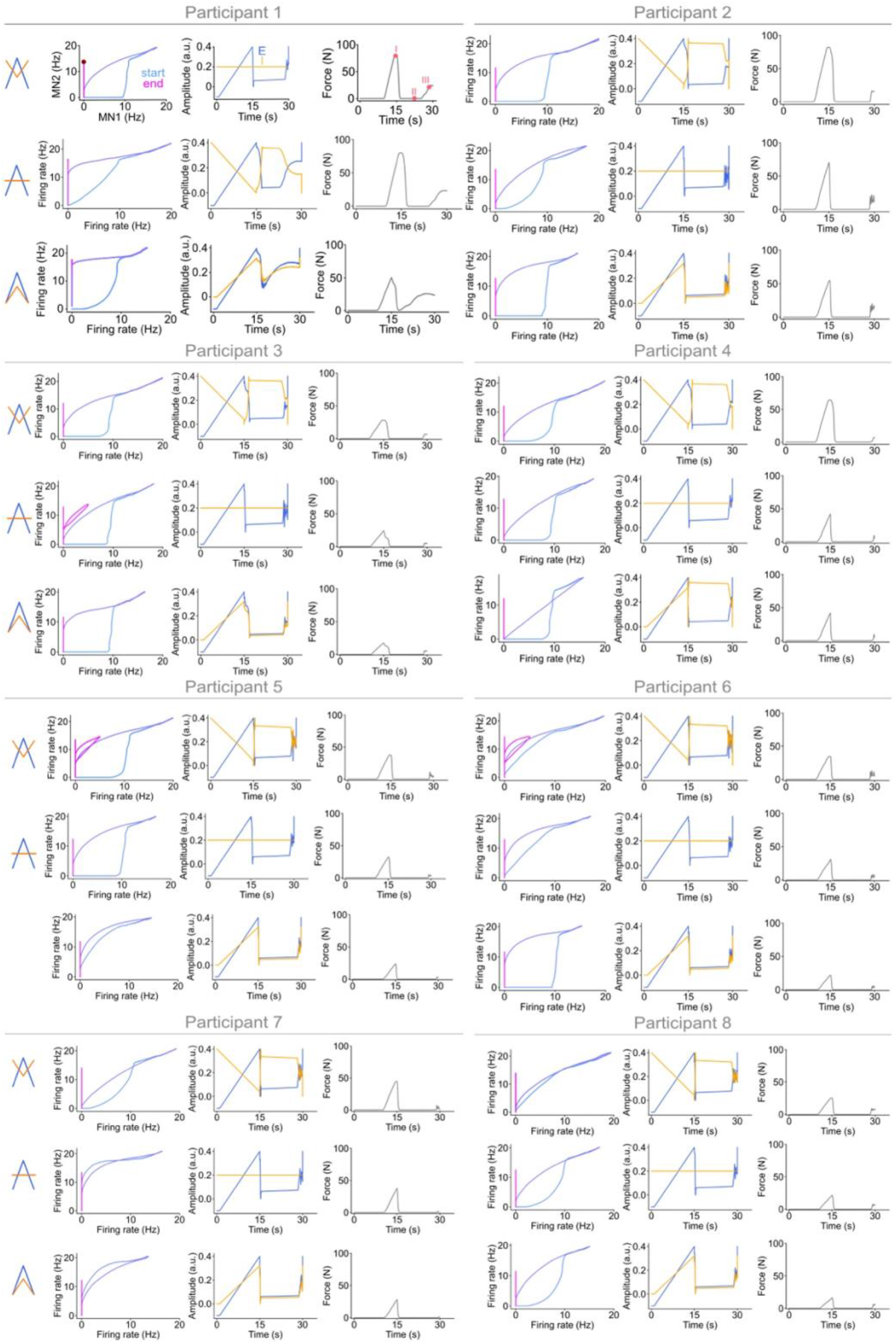
Simulated cursor-control task for another five participants. The following motoneuron pairs are shown: Participant 1, motoneuron pair 45-51 (Δ*V*_*thCa*_ ~ 3), Participant 2, motoneuron pair 29–33 (Δ*V*_*thCa*_ ~ 3); Participant 3, motoneuron pair 2–5 (Δ*V*_*thCa*_ = 3.7); Participant 4, motoneuron pair 25–30 (Δ*V*_*thCa*_ = 2.3); Participant 5, motoneuron pair 29–31 (Δ*V*_*thCa*_ = 4.2); Participant 6, motoneuron pair 24–26 (Δ*V*_*thCa*_ = 4.6); and Participant 7, motoneuron pair 45–47 (Δ*V*_*thCa*_ = 2.3). Data presented as in Figure 4d and Suppl. Figure 6. Note that force traces displayed a characteristic end-of-trial increase, closely mirroring the experimental data. Data presented as in Figure 4d.

**Supplementary Fig. 5.**
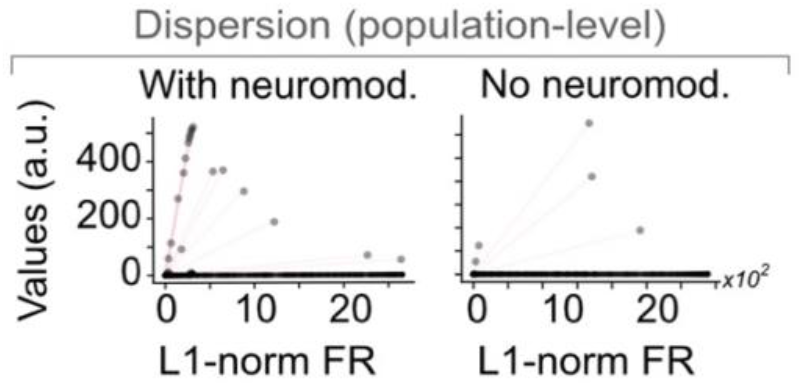
Dispersion for the online cursor-control task over the whole trial. Example data for Participant 1.

**Supplementary Fig. 6.**
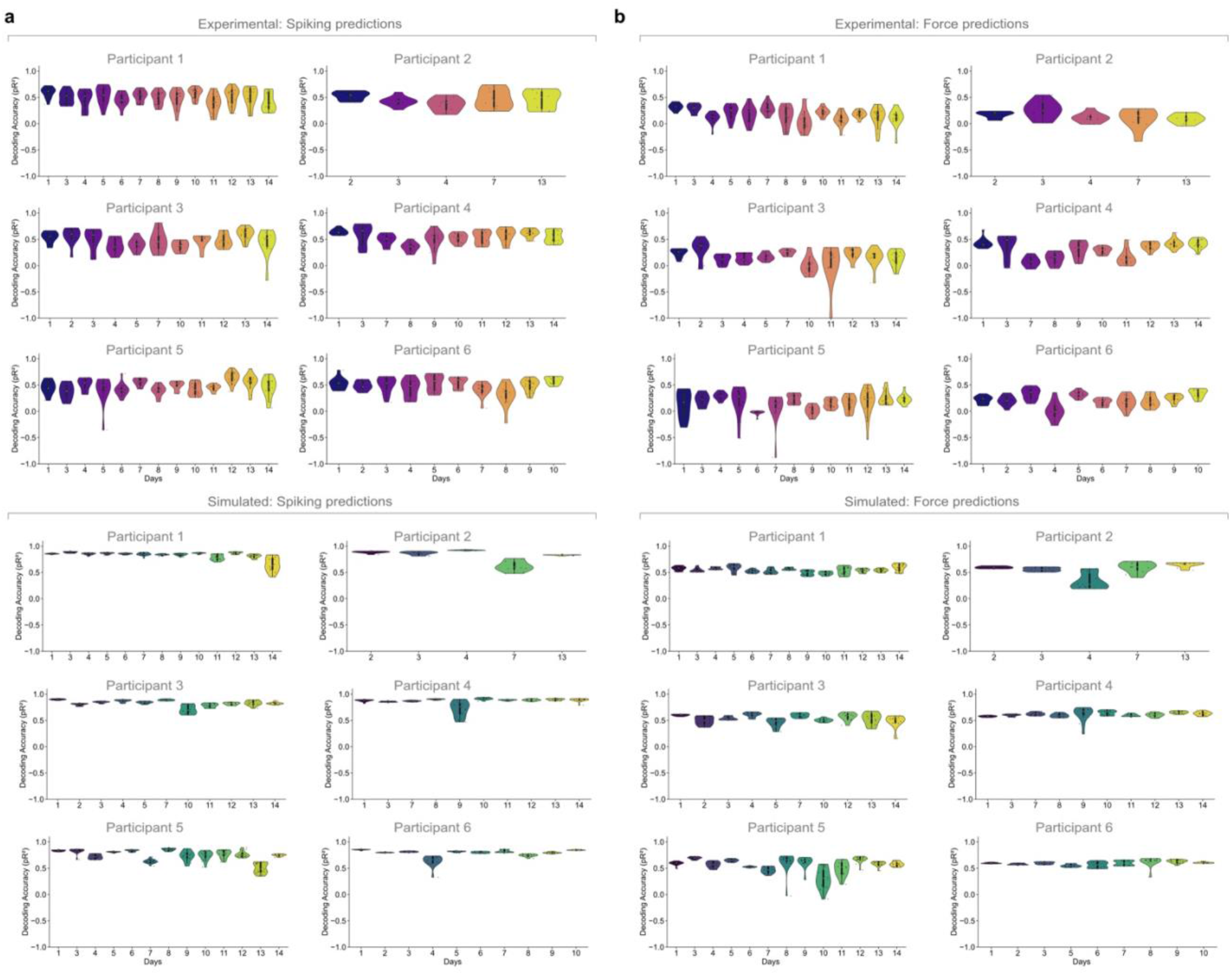
Poisson GLM analyses to predict the spiking of each motoneuron from the spiking of the other recorded motoneurons (a) or the measured force (b). Violin, distribution for each session; markers, individual predictions. Top plots, experimental data; bottom plots, simulated data.

**Supplementary Fig. 7.**
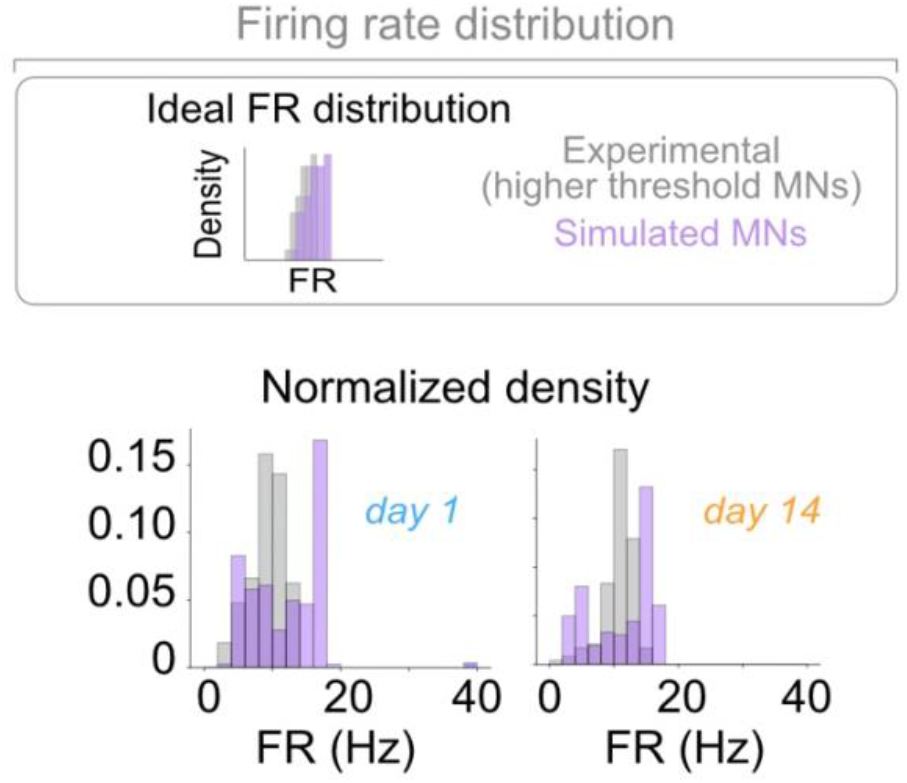
Firing rate distributions for experimental and simulated cursor control data. Firing-rate distributions were matched between experimental and simulated data, with simulations requiring a bias toward higher firing rates to capture the activity of high-threshold motoneurons observed experimentally. Data for Participant 1.

**Supplementary Table 1.** Morphological and passive membrane parameters.

| Variable | Definition | Value |
| --- | --- | --- |
| $\rho$ | Intracellular resistivity | 70 $\Omega \cdot \text{cm}$ |
| $c_m$ | Membrane capacitance | 1 $\mu\text{F} \cdot \text{cm}^{-2}$ |
| $r_s$ | Soma radius | 0.003875–<br>0.004125 cm |
| $l_s$ | Soma length | 0.00775–0.00825<br>cm |
| $r_d$ | Dendrite radius | 0.002075–<br>0.003125 cm |
| $l_d$ | Dendrite length | 0.55–0.68 cm |

